# Heterogeneous evolution of effective recombination landscapes across percomorph fishes

**DOI:** 10.64898/2026.09.28.754948

**Authors:** Marion Talbi, Marius Roesti, Catherine L. Peichel, Martin Reichard, Diego Hartasánchez, Thibault Latrille, Nicolas Salamin, Walter Salzburger, Stephan Peischl, Ole Seehausen, Milan Malinsky

**Affiliations:** Division of Aquatic Ecology and Evolution, Institute of Ecology and Evolution, University of Bern, Switzerland; Department of Fish Ecology and Evolution, EAWAG, Kastanienbaum 6047, Switzerland; Division of Evolutionary Ecology, Institute of Ecology and Evolution, University of Bern, Switzerland; Institute of Vertebrate Biology, Czech Academy of Sciences, Brno, Czech Republic; Department of Botany and Zoology, Masaryk University, Brno, Czech Republic; Department of Ecology and Vertebrate Zoology, University of Lodz, Lodz, Poland; Department of Computational Biology, Génopode, University of Lausanne, Switzerland; Center for Research and Advanced Studies of the National Polytechnic Institute, Mexico City, Mexico; Zoological Institute, Department of Environmental Sciences, University of Basel, Basel, Switzerland; Interfaculty Bioinformatics Unit, University of Bern, Switzerland

## Abstract

Recombination shapes allelic combinations across the genome and plays a central role in adaptation and speciation. While recombination rates are known to vary and evolve, how recombination landscapes diverge between populations and which processes drive this divergence remains poorly understood. Here, we use the Population Recombination Divergence Index (PRDI) to quantify divergence in effective recombination landscapes between seven pairs of populations and species from four percomorph fish lineages (cichlids, sticklebacks, killifish, and clownfish). While accounting for stochasticity and confounders of linkage disequilibrium (LD) -based recombination rate estimates, we find that recombination landscapes diverged substantially in all comparisons. This divergence generally increases with genetic differentiation and divergence time, but the amount of PRDI accumulated per unit of genetic divergence (F_ST_), net nucleotide divergence (d_a_), or divergence time varies markedly among population and species pairs. Comparisons across systems suggest that PRDI accumulation tends to be faster (per unit of F_ST_, d_a_, or divergence time) in pairs with higher inferred gene flow. We also introduce a local version of PRDI (*PRDI_l_*) to better characterize genomic heterogeneity in effective recombination divergence. We find that higher inferred gene flow between populations tends to be associated with stronger intragenomic correlation between local F_ST_ and *PRDI_l_*. Finally, we propose a conceptual framework for investigating how divergent selection and gene flow may interact to shape effective recombination landscape divergence. Our work highlights the heterogeneous dynamics of effective recombination landscape evolution, and motivates further theoretical and empirical efforts to integrate recombination landscape evolution into studies of adaptation and speciation.

## Introduction

Meiotic recombination – the reciprocal exchange of genetic material between chromosomes during meiosis – is universal among sexually reproducing species (Barton and Charlesworth 1998; Barton et al. 2007; Stapley et al. 2017). It plays a crucial mechanistic role in ensuring proper pairing and segregation of homologous chromosomes (Handel and Schimenti 2010; Capilla et al. 2017). Beyond this role in meiosis, recombination also has major evolutionary consequences. Each generation, it reshuffles genetic variation, breaking down existing haplotypes and generating new ones that contain novel, often unique, combinations of genetic variants (Barton et al. 2007; Barton 2022).

By generating novel haplotypes, recombination can facilitate rapid adaptation by enabling selection to act more effectively on standing genetic variation, rather than relying solely on rare *de novo* mutations (Hermisson and Pennings 2005; Barrett and Schluter 2008). In some cases, this can lead to extremely rapid adaptation or combinatorial speciation (Marques et al. 2019; Rudman et al. 2022; Meier et al. 2023). This effect of bringing together beneficial variants and facilitating the purging of deleterious variants by decoupling them from their background is considered the leading explanation for the existence and prevalence of sex and recombination across the tree of life (Felsenstein 1974; Barton and Charlesworth 1998; Otto and Lenormand 2002; Nielsen 2006). However, recombination can also hinder adaptation. During adaptive divergence with gene flow – that is, when populations experience divergent selection but are not fully reproductively isolated – migration and recombination counteract divergent selection. While selection within each population acts to couple alleles that are beneficial in each population’s niche, gene flow and recombination continually break apart these advantageous combinations. Thus, adaptive population divergence proceeds only if selection is sufficiently strong or if recombination among the involved loci is reduced (Felsenstein 1981; Barton and Bengtsson 1986; Feder et al. 2012; Aeschbacher et al. 2017; Berner and Roesti 2017; Samuk et al. 2017).

Recombination rates are not static – they evolve (Samuk et al. 2020). Recombination rates are highly variable both between and within species (Stapley et al. 2017; Henderson and Bomblies 2021). This variation affects not only the overall rate of recombination, but also its intragenomic distribution, i.e., the recombination landscape, at both broad and fine scales (Myers et al. 2005; Auton and McVean 2012). To understand how and why recombination landscapes evolve, it is important to quantify recombination landscape evolution across a range of natural populations and related species (Ortiz-Barrientos et al. 2016; Ortiz-Barrientos and James 2017; Ravinet et al. 2017). Despite considerable efforts, accurate inference of recombination landscapes from genomic data remains challenging (Peñalba and Wolf 2020). Pedigree-based approaches provide valuable insights (Halldorsson et al. 2019; Kivioja and Rastas 2024; Venu et al. 2024); however, they typically yield limited resolution and require genetic data from large families, which are difficult and sometimes impossible to obtain for natural populations (Peñalba and Wolf 2020). Gamete-based approaches (Hinch et al. 2019; Sun et al. 2019; Xu et al. 2019) are also often limited in resolution (but see (Malinsky et al. 2025)). They can be applied across a broader range of species and provide a direct snapshot of contemporary crossover events, but require thousands of gametes, which means that only male gamete-based maps have been produced so far, which may differ considerably from female maps, and therefore not be representative of the true recombination distribution within the population (Sardell and Kirkpatrick 2020).

In contrast, linkage disequilibrium (LD)-based approaches provide very high resolution and reflect recombination events accumulated across hundreds of thousands of generations, effectively averaging male and female rates. With the decreasing cost of whole-genome sequencing, LD-based maps have become relatively straightforward to infer across a broad range of organisms (Myers et al. 2005; Auton and McVean 2012; Chan et al. 2012; Singhal et al. 2015; Terhorst et al. 2017; Beeson et al. 2019; Spence and Song 2019; Déserts et al. 2021; Hoge et al. 2024). In interpreting the results, it is crucial to take into account that LD-based maps reflect the population recombination rate (*ρ*), which is the product of the recombination rate (r) and effective population size (*N_e_*); specifically, *ρ* = 4 ∗ *N_e_* ∗ *r* (Chan et al. 2012; Terhorst et al. 2017; Spence and Song 2019). Meaningful quantitative comparisons of population recombination landscapes across populations or species have been difficult because of randomness (sampling, drift), methodological noise, and the effects of processes such as demographic changes or selection on linked loci, which affect *ρ* via genome-wide or local changes in *N_e_*. To overcome these challenges, we recently developed the Population Recombination Divergence Index (PRDI), a metric that allows robust comparison of LD-based recombination maps by explicitly accounting for statistical variance and multiple confounding factors (**Fig. 1**) (Talbi et al. 2025).

**Figure 1:**
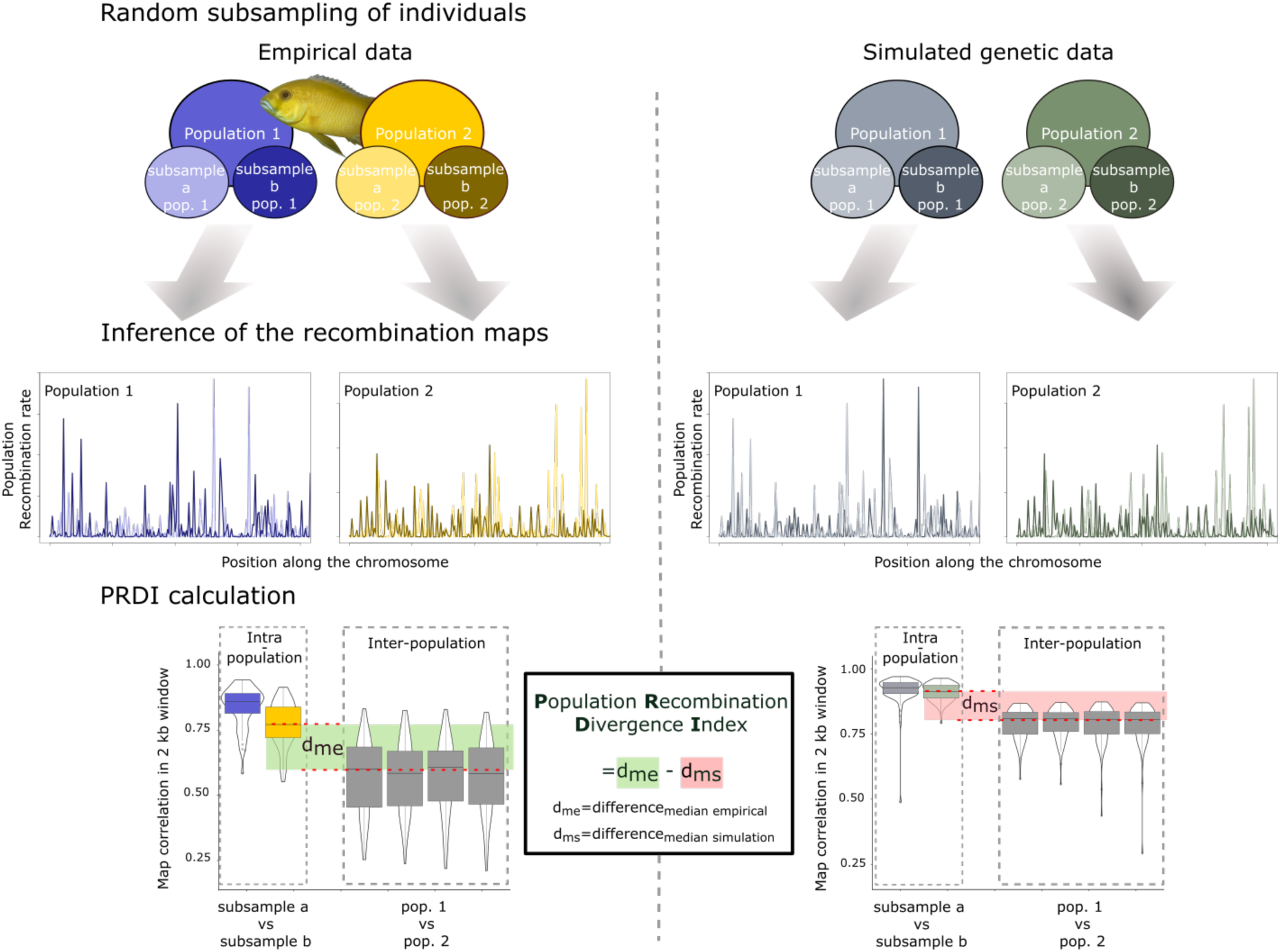
The methodology behind the Population Recombination Divergence Index (PRDI). We developed the PRDI to facilitate recombination map comparisons while controlling for two factors: (1) sensitivity to sampling variance and methodological noise, and (2) differences in demographic (*N*_e_) histories between populations. See Talbi et al. (2025) for details. **Random subsampling of individuals**: To quantify the impact of sampling variability, we divided the sampled individuals from each population into two independent subsets. **Inference of the recombination maps:** We reconstructed a recombination map from each subset. The differences between the two maps inferred from subsets from the same population reflect the amount of sampling variance and methodological noise. **PRDI calculation:** We calculated correlations between the inferred maps by grouping the comparisons into two categories: intra-population comparisons (subsets from the same population), and inter-population comparisons (population 1 vs. population 2). By subtracting the median inter-population correlation from the median intra-population correlation, we obtained *d_me_*, the median empirical difference. **Right panel:** To control for differences in demographic histories between the two populations that could affect LD and therefore the inferred maps, we repeated the same procedure on simulated genetic data to obtain *d_ms_*. These data were generated using the evolutionary parameters inferred from the focal populations. **PRDI calculation (middle)**: We calculated PRDI as the difference between *d_me_* and *d_ms_*.

In Talbi et al., we applied the PRDI framework to a single pair of populations of the cichlid fish *Astatotilapia calliptera,* representing recent adaptive divergence with substantial ongoing gene flow along a depth gradient in a small crater lake. Here, we extended this approach to quantify recombination landscape divergence across seven comparisons involving 14 populations from four percomorph fish lineages that differ in ecology, evolutionary history, and levels of gene flow: cichlids (including the two Lake Masoko populations from Talbi et al. (2025)), stickleback, clownfish, and killifish. First, we used PRDI to quantify recombination landscape divergence across the comparisons and found that recombination landscape divergence tends to scale with genetic differentiation, sequence divergence, and time since divergence from their common ancestor. Second, we found substantial heterogeneity in the rate of recombination landscape divergence across comparisons. As we knew that gene flow rates vary among our populations and species pairs, we investigated whether variation in gene flow could contribute to these rate differences. Finally, we developed and applied a local version of PRDI (*PRDI_l_*) to quantify intragenomic heterogeneity in recombination landscape divergence and examined how gene flow, genetic divergence (F_ST_), and recombination landscape divergence interact locally within the genome. This study provides a comparative empirical assessment of recombination landscape evolution across wild populations of teleosts and provides new tools and conceptual insights into the study of recombination evolution.

## Results

### Sampling and data for population comparisons

We generated ∼15x whole-genome resequencing data for 637 individuals and, after aligning to chromosome-level genome assemblies, produced single nucleotide polymorphism (SNP) datasets for the four groups (**Fig. 2**; Methods; **STable 1**): (i) cichlids (282 individuals, six populations, 31.5 million SNPs), (ii) stickleback (172 individuals, four populations, 13.0 million SNPs), (iii) killifish (83 individuals, two populations, 15.8 million SNPs), and (iv) clownfish (85 individuals, two populations, 27.1 million SNPs). These datasets formed the basis for all subsequent analyses.

**Figure 2:**
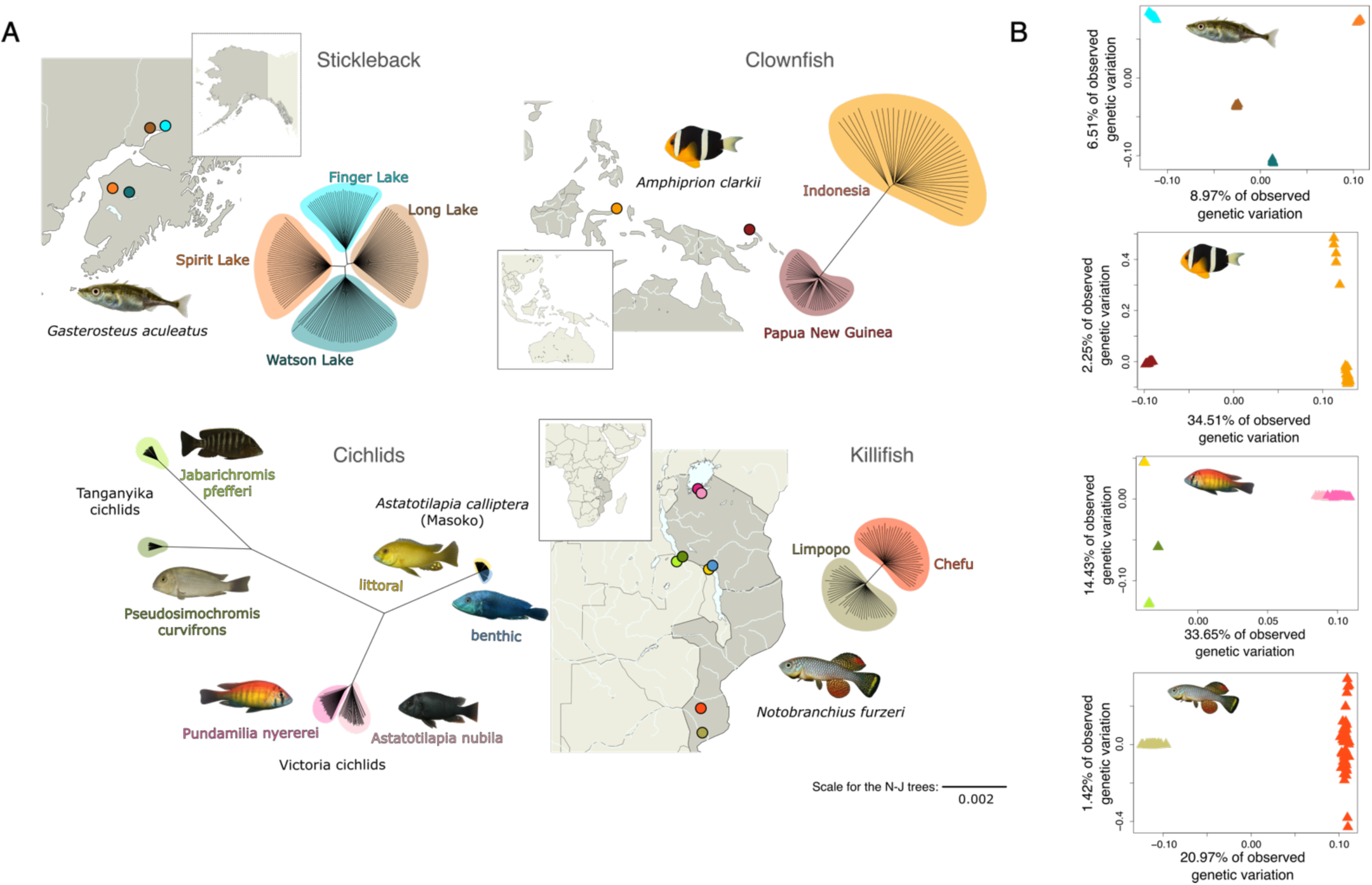
Maps and global genetic structure for each dataset. **(A) Top left**: The stickleback dataset includes four populations from allopatric lakes in Alaska: Finger Lake and Long Lake in the north, and Watson Lake and Spirit Lake in the south. Blue indicates benthic populations, while orange/brown indicates limnetic populations. **Top right**: The clownfish dataset includes two populations of *Amphiprion clarkii* from Papua New Guinea and Indonesia. **Bottom left:** The cichlid dataset is composed of six populations from three main radiations. The exception is *A. calliptera* from Lake Masoko, a crater lake north of Lake Malawi, where littoral and benthic populations represent two ecotypes of the same species. *P. nyererei* and *A. nubila* are from Lake Victoria, with *A. nubila* representing one of the ancestral lineages that contributed to the hybrid origin of the radiation from which *P. nyererei* diversified. *J. pfefferi* and *P. curvifrons* are from Lake Tanganyika and diverged millions of years ago. **Bottom right:** The killifish dataset includes two populations of *Nothobranchius furzeri* from distinct catchment basins in Mozambique: Chefu and Limpopo rivers. The cichlid and killifish populations are both represented on the bottom map. Neighbor-joining trees are shown on the same scale. (**B**) Principal component analyses showing the genetic structure for each dataset.

We started by examining population genetic structure within each system using principal component analyses (PCA) and neighbor-joining (NJ) trees to select individuals while avoiding strongly structured samples that could confound downstream analyses. Overall, we found that most of the populations were genetically homogeneous, with no sub-structure visible either in the PCAs or in the NJ-trees. A small degree of sub-structure was apparent in one clownfish and in one killifish population (**Fig. 2**). Given that this sub-structure is weak relative to the between-population variation within each system, as reflected by the small within-population distances on the NJ trees, we decided to keep these populations as such for all following analyses.

Genetic diversity is a major factor in determining the accuracy of LD-based recombination rate inference, because these methods rely on LD patterns among SNPs within populations (Raynaud et al. 2023). Therefore, for PRDI comparisons, we selected pairs of populations with similar levels of genetic diversity (**Table 1**). We also selected the pairs for comparisons to be sister populations or species in our datasets. Therefore, despite multiple comparisons within some groups (cichlids, stickleback), the comparisons are phylogenetically independent. To account for differences in demographic history that can affect LD-based recombination inference, we reconstructed past effective population size (*N_e_*) history for each population using smc++ (**SFig. 1**, Terhorst et al. 2017). These histories were used as demographic inputs in pyrho (Spence and Song 2019) to model the expected LD patterns under each population’s inferred demographic history and estimate population-specific recombination rates.

**Table 1:**
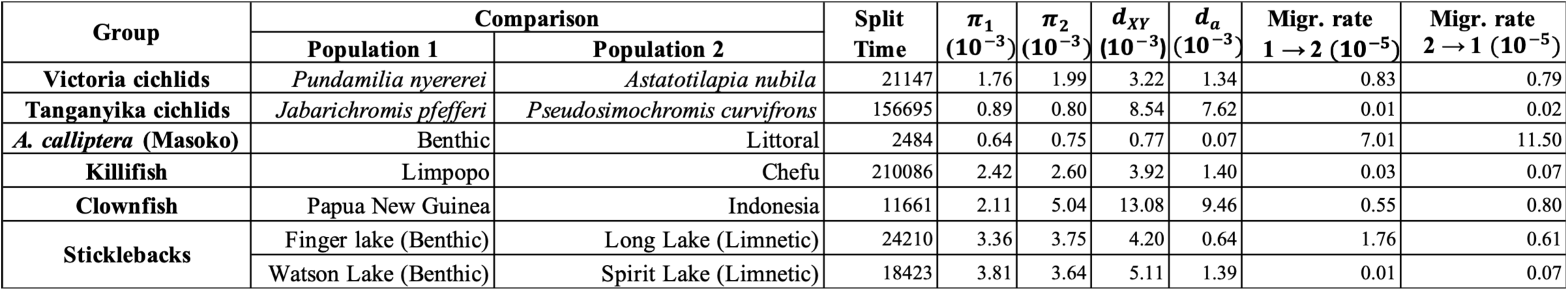
Population comparisons with key demographic and genetic information. For each pair of populations/species that we compare, we calculated: 1) mean genetic diversity for each populations (*π*_1_ and *π*_2_); 2) absolute nucleotide divergence (*d_XY_*), and net nucleotide divergence (*d_a_*); 3) split time estimate obtained with smc++; 4) migration rate estimates (a measure of gene flow) obtained with fsc2.

Although pyrho reports its estimates in units of r, these estimates are LD-based. The pattern of LD from which they are inferred reflects the combined effect of the underlying crossover distribution and population-level evolutionary processes such as selection, migration, and drift. Pyrho obtains its estimates by converting the population recombination parameter *ρ* using a genome-wide estimate of *N_e_*. Its estimate cannot therefore be interpreted as either a direct measurement of r or of *ρ*. We thus refer in the rest of the manuscript to pyrho estimates using the more general term ‘effective recombination rates’ (*r_e_*), with ‘effective’ referring to the fact that they carry only signatures of recombination that are transmitted through generations across the genealogical ancestry of the samples.

The selected pairs have a range of estimated split times from smc++ between 2,500 and 210,000 generations ago (**Table 1**; **SFig. 1**), and gene flow estimates from fastsimcoal2 varying over three orders of magnitude from 1e-07 to 1.15e-04 (**Table 1**; Methods). For most pairs, split time estimates were consistent with estimates from the literature (Bartáková et al. 2013; Malinsky et al. 2015; Härer et al. 2021; Marques et al. 2025). However, split times for the Tanganyika cichlids and the clownfish pairs were considerably lower than literature-based estimates (Tanganyika: 150,000 vs 900,000 generations in Ronco et al. (2021); clownfish: 11,000 vs 60,000 generations in Schmid et al. (2024). This is likely because smc++ assumes no gene flow after divergence. We thus decided to keep the split times from smc++ for all comparisons except the clownfish and Tanganyika cichlids, for which we used estimates from the literature (i.e., 60,000 and 900,000 generations respectively). To characterize genetic diversity within and among population pairs, we calculated several standard population genetic statistics, including nucleotide diversity (*π*), absolute genetic divergence (d_xy_), and net nucleotide divergence (d_a_; (Cruickshank and Hahn 2014)). The mean values are shown in **Table 1**.

### Recombination map characterization

The reconstructed maps returned by pyrho varied substantially in their general characteristics. Although recombination hotspots are present in all maps, we found that overall, recombination events are distributed more broadly across the genome than in the human and bird (zebra finch) maps used for comparisons (**Fig. 3A**, zebrafinch and human from (Singhal et al. 2015; Halldorsson et al. 2019); **SFig. 2**). This difference in comparison to human maps is expected; in human populations, recombination hotspots are directed by PRDM9 and are expected to be very prominent (Myers et al. 2005, 2010; Baudat et al. 2010), whereas percomorph fishes lack this PRDM9-directed mechanism (Baker et al. 2017). However, the differences we observed in comparison to the zebra finch, which also lacks the PRDM9 mechanism (Singhal et al., 2015), are notable. Especially in the killifish populations, recombination events are distributed much more broadly across the genome, with 50% of recombination events concentrated within 25% of the sequence as compared to ∼8% in zebra finch (Singhal et al. 2015).

**Figure 3:**
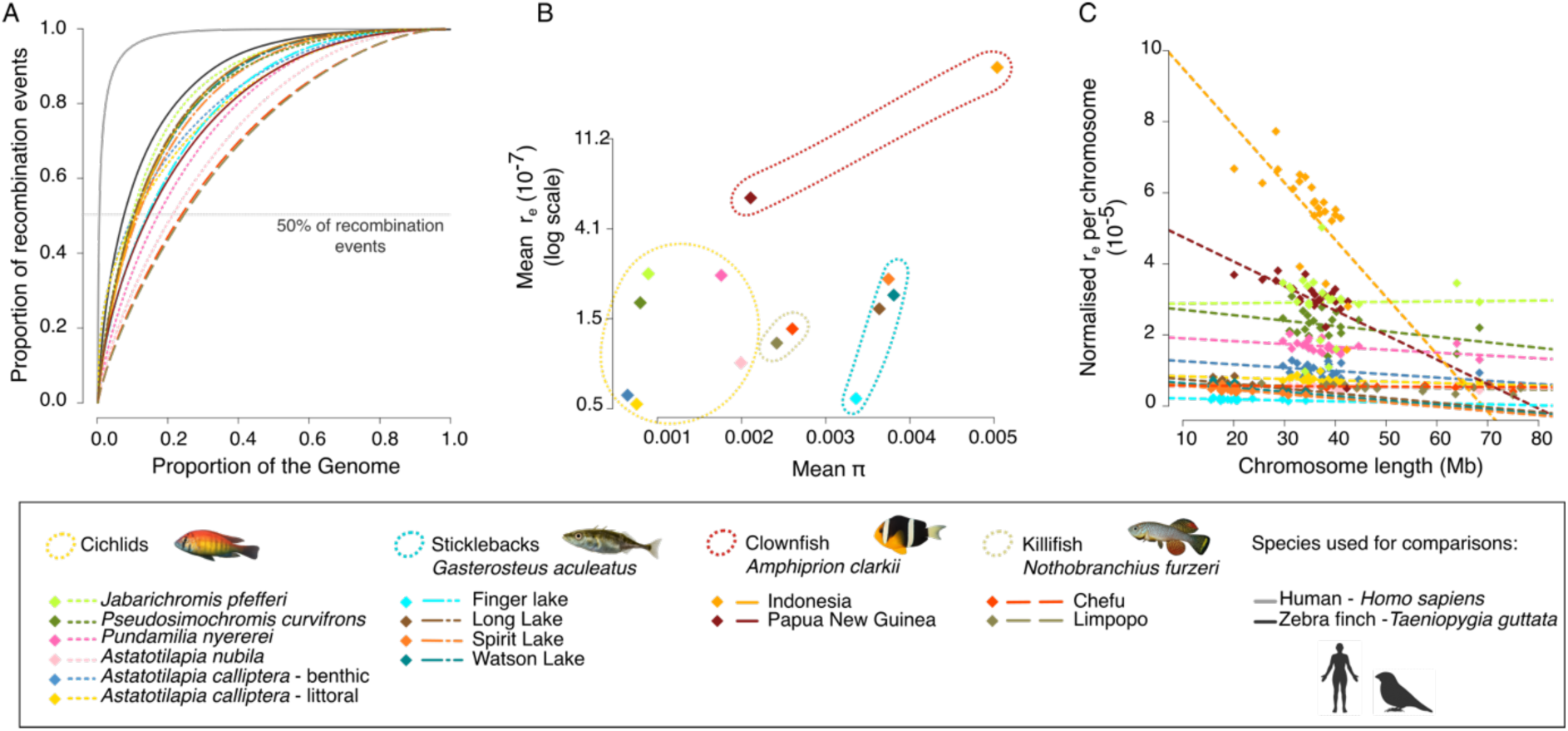
Recombination rate distribution and relationship with *π*. **(A)** Cumulative curves representing the fraction of the genome needed to account for a given proportion of inferred recombination events for each of the 14 populations, including two other species, *Homo sapiens* and *Taeniopygia guttata*, in grey and black, respectively, for comparison. The horizontal line corresponds to 50% of recombination events. **(B)** The positive relationship between the level of genetic diversity, *π*, and the effective recombination rate, *r_e_*. **(C)** To control for differences in diversity between populations, we normalised *r_e_* per *π*, and plotted this value against the chromosome length.

Overall, Masoko cichlids have the lowest recombination rates, with a median *r_e_* value of 3.29×10^-9^. Tanganyika and Victoria cichlids have very similar medians (*r*_e_ = 7.27×10^-9^ and *r*_e_ = 9.94×10^-9^, respectively), as with sticklebacks (*r*_e_ = 6.25×10^-9^) and killifish (*r*_e_ = 1.03×10^-8^). Finally, clownfish showed a median *r*_e_ of 4.94×10^-8^, roughly an order of magnitude higher than the others (**Fig. 3B**; **SFig. 3**). Because the reduction in genetic diversity due to linked selection depends on recombination rate (Corbett-Detig et al. 2015; Elyashiv et al. 2016), and because recombination itself is mutagenic (Halldorsson et al. 2019), we examined the relationship between *π* and *r_e_*. Across populations, *π* and *r_e_* showed a positive correlation, although this was not significant (**Fig. 3B**; Pearson *ρ* = 0.51, p-value = 0.059; Spearman *ρ* = 0.37, p = 0.19).

The number of crossovers per chromosome is known to be constrained from a minimum of one crossover per chromosome, needed for proper meiotic segregation, to a maximum of not much more than one, due to crossover interference – non-independence of crossovers (Otto and Payseur 2019). These constraints predict higher recombination rates on smaller chromosomes (Haenel et al. 2018). Consistent with this expectation, we observed overall a negative association between *r_e_* and chromosome size. Interestingly, the strength of this relationship varied greatly among groups, suggesting considerable variability despite the constraints (**Fig. 3C**; estimates and p-values in **Table S2**).

### Scaled recombination divergence varies across comparisons

To quantify the degree of recombination landscape divergence between populations or species, we calculated the Population Recombination Divergence Index (PRDI; **Fig. 1**; Methods; Talbi et al., 2025) for each taxon pair. PRDI was positive across all seven comparisons and varied from 0.03 in the Victoria cichlid pair to 0.29 in the Tanganyika cichlid pair (**Table S3**), which indicates more effective recombination landscape divergence than expected if the underlying crossover landscapes remained the same – while accounting for demographic changes, methodological noise (e.g., variant calling errors), and sampling variance. These results indicate either divergence of crossover landscapes, selection-driven variation in effective recombination landscapes, or a combination of the two.

As it is currently unknown how and why effective recombination landscape divergence accumulates, we tested, across comparisons, the extent to which PRDI scales with genetic differentiation (F_ST_), with net nucleotide divergence (*d_a_*), and with the time since the split from the common ancestor. We found a tendency for positive rank associations in all three comparisons, albeit none was significant, given the low statistical power afforded by seven comparisons: PRDI and F_ST_ (Spearman *ρ* = 0.61, p-value = 0.16; **Fig. 4A**), PRDI and *d_a_*(Spearman *ρ* = 0.5; p = 0.27, **SFig. 4A**), and PRDI with divergence time (Spearman *ρ* = 0.54; p = 0.24, **SFig. 5A**). These results suggest that recombination landscape divergence tends to increase with genetic divergence in allele frequencies (F_ST_), net nucleotide divergence, and divergence time. The trends became weaker but remained positive after excluding the *J. pfefferi* vs *P. curvifrons* cichlid comparison which could be considered an outlier (F_ST_: Spearman *ρ* = 0.37, p = 0.49, **Fig. 4B**; d_a_: Spearman *ρ* = 0.26, p = 0.65, **SFig. 4B**; Split time: Spearman *ρ* = 0.26, p = 0.66, **SFig. 5B**).

**Figure 4:**
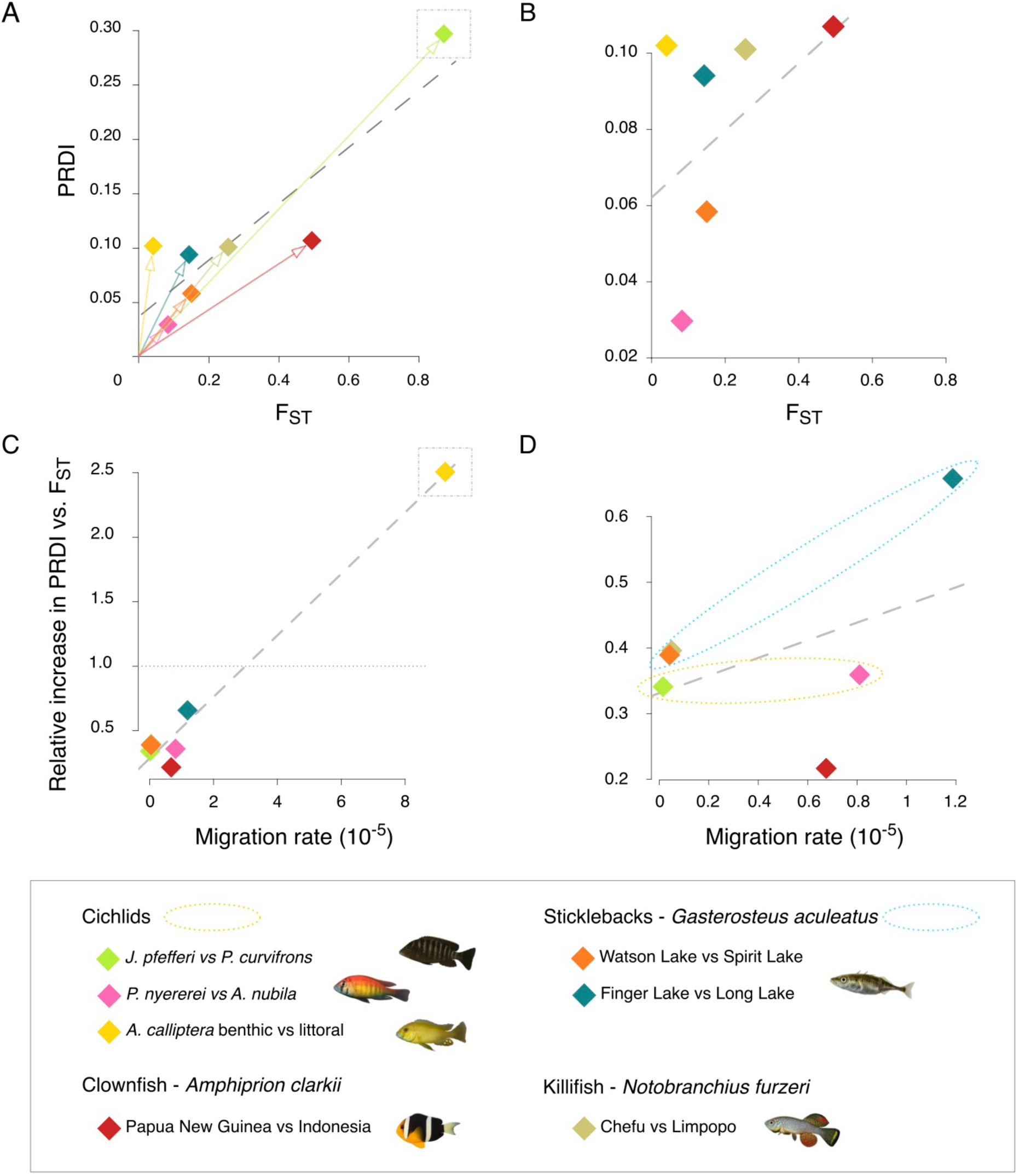
Recombination landscape divergence increases with genetic differentiation and gene flow. **(A)** Relationship between the population recombination divergence index (PRDI) and mean genome-wide genetic differentiation (F_ST_) for each population pair. Panel (**B**) shows the same relationship without the *J. pfefferi* and *P. curvifrons* comparisons (light green diamond in the dotted rectangle in panel A). Each coloured symbol represents one pair. Arrows from the origin highlight that the slope, representing PRDI per unit of divergence, differs among comparisons. **(C)** Relationship between the scaled recombination landscape divergence, calculated as PRDI divided by mean F_ST_, and migration rate. The dotted horizontal line indicates PRDI/F_ST_ equal to 1 as a visual reference. (**D**) Same relationship without the *A. calliptera* comparison (yellow diamond in the dotted rectangle in panel C). Dotted ellipses highlight the patterns within stickleback (blue) and cichlid (yellow) systems.

**Figure 5:**
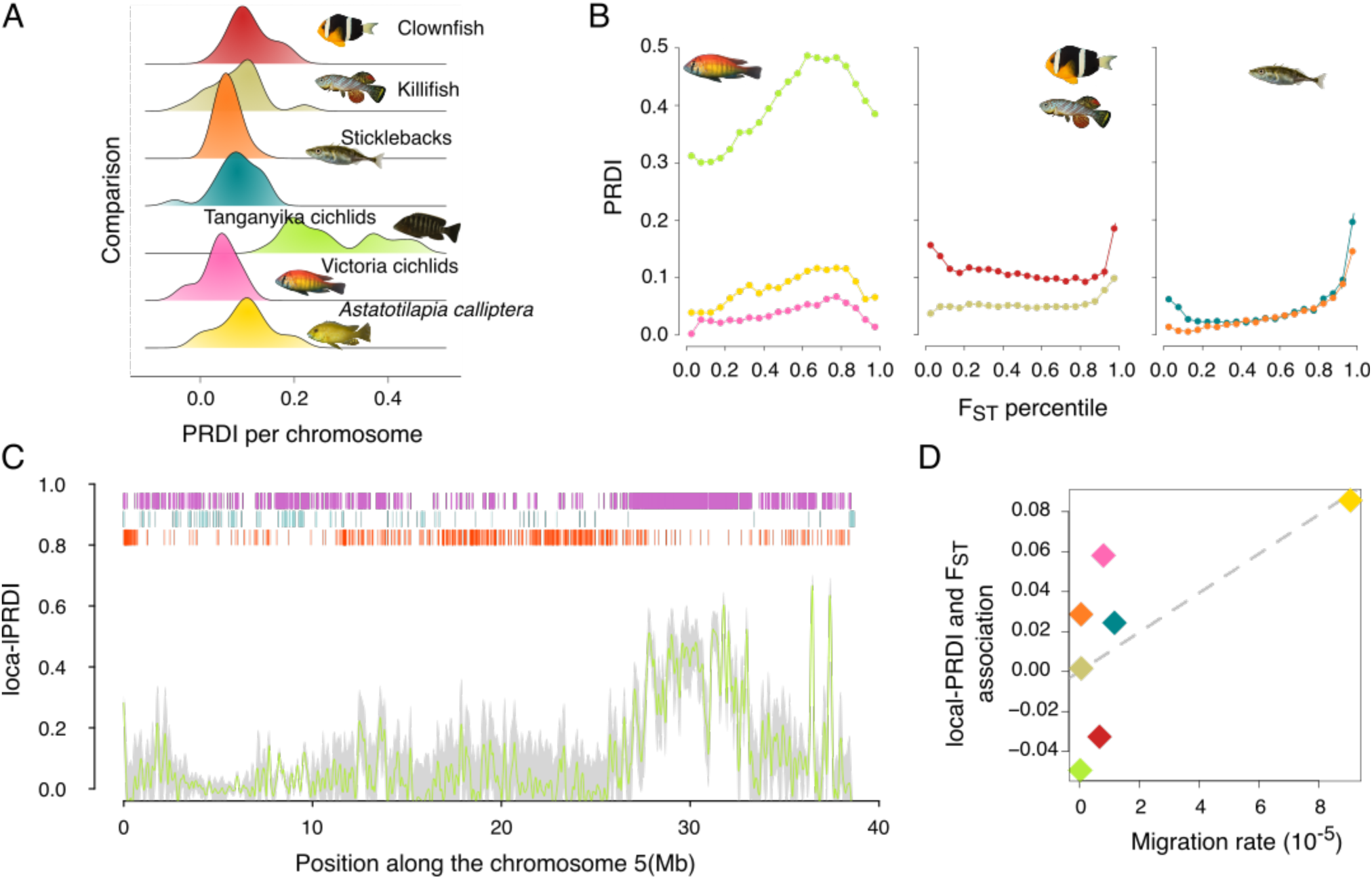
Heterogeneity in recombination landscape divergence and local-PRDI. **(A)** Distribution of chromosome-level PRDI for each population pair. **(B)** Relationship between PRDI and F_ST_ within each comparison. For each pair, the genome was split into F_ST_ percentiles, and PRDI was calculated for each percentile bin. The lowest percentile corresponds to the lowest F_ST_ values. **(C)** Example of local PRDI along one chromosome of the *J. pfefferi* and *P. curvifrons* comparison. The green line shows local PRDI in 2kb windows, and the grey area shows a confidence interval based on 3 standard deviations. Bars at the top indicate genomic regions with strongly diverged recombination in purple (i.e., when *dm_e_* > *dm_s_* + 5 ∗ *sd_dms_*) and conserved recombination in light blue (i.e., when *dm_e_* < *dm_s_* + 5 ∗ *sd_dms_*). Red bars represent 5% F_ST_ outlier windows. **(D)** Strength of the association between local PRDI and F_ST_ (Spearman correlation in 50kb windows) plotted against migration rate. Each point represents a population pair, coloured as in Figure 4.

Despite the overall tendency for PRDI to increase with genetic divergence, we found that the amount of effective recombination divergence accumulated per F_ST_ unit varied substantially across the different comparisons (as indicated by the arrows in **Fig. 4A, SFig. 4A**, and **SFig. 5A**). This raises the question of which additional processes may contribute to this heterogeneity. In diverging populations, gene flow and divergent selection can jointly reshape patterns of linkage disequilibrium by continually introducing and removing ancestry blocks and by altering the persistence of locally advantageous or deleterious allele combinations. These processes may generate differences in effective recombination landscapes even without changes in the underlying crossover distribution, although selection on crossover rates or genomic architecture may also contribute (Bank et al. 2012; Ortiz-Barrientos et al. 2016; Kirkpatrick and Barton 2006). We therefore tested whether the inferred magnitude of gene flow between sister taxa was associated with the amount of recombination landscape divergence accumulated per unit of genetic divergence or divergence time.

Although not statistically significant, all associations between the scaled PRDI and migration rate were positive ((1) F_ST_: Spearman *ρ* = 0.61, p-value = 0.17, **Fig. 4C**; (2) d_a_: Spearman *ρ* = 0.46, p-value = 0.3, **SFig. 4C**; (3) divergence time: Spearman *ρ* = 0.75, p-value = 0.06; **SFig. 5C**). The *A. calliptera* benthic vs littoral comparison in cichlids (in yellow on **Fig. 4**) has high values of scaled PRDI (high rates of recombination landscape divergence) and can be considered an outlier. Excluding this comparison reduced the strength of the trends; nevertheless, all trends remained positive ((1) Spearman *ρ* = 0.14, p-value =0.8; (2) Spearman *ρ* = 0.37, p-value = 0.49; (3) Spearman *ρ* = 0.6, p-value =0.24; **Fig. 4D**, **SFig. 4D, SFig. 5D**) and were also observed within lineages (i.e., within cichlids and sticklebacks; see dotted circles on **Fig. 4D**).

### Local PRDI captures the heterogeneity of effective recombination divergence along the genome

To investigate how recombination landscape divergence varies across the genome, we first separated the PRDI measures by chromosome. We observed considerable variation in almost all comparisons, with effective recombination landscapes showing either more or less divergence than expected from demographic changes, genetic drift, and sampling variance alone (**Figure 5A**). The number of chromosomes with significantly greater effective recombination landscape divergence than expected ranged from two of 22 chromosomes for the Victoria cichlid comparison to 18 of 20 for the Spirit-Watson stickleback comparison, and up to all chromosomes for the Tanganyika cichlid and the clownfish comparisons. There were a small number of chromosomes with a negative PRDI, indicating less divergence than expected. Specifically, we observed this in *A. calliptera* from Masoko, Victoria cichlid, killifish, and Finger-Long stickleback comparisons, which all had one or a few chromosomes with less recombination divergence than expected. However, none of these estimates were significantly below zero. The Tanganyika cichlid comparison stands out with a distinct bimodal distribution, with eight chromosomes showing PRDI values approximately 20% higher than the other chromosomes. This pattern may reflect chromosomal rearrangements segregating between the two species, as reported previously in other Tanganyika cichlids (Fischer et al. 2021). An exhaustive list of chromosome-level PRDI values and their associated 95% confidence intervals is provided in **Table S4**.

To investigate the factors that contribute to this variation in PRDI, we next grouped genomic regions according to percentiles of F_ST_ and calculated PRDI within each 5% interval (e.g., 0 to the 5^th^ percentile of F_ST_, then the 5^th^ to the 10^th^ percentile of F_ST_, etc.). This allowed us to assess, in aggregate across the genome, the relationship between PRDI and F_ST_. The strength and shape of this relationship varied notably among the comparisons, with three distinct patterns: 1) in sticklebacks, PRDI tends to increase with F_ST_, despite being also higher at low F_ST_ in one pair; 2) in cichlids, the relationship increases up to intermediate levels of divergence before decreasing at higher values; 3) in killifish and clownfish, the relationship is much flatter, and even slightly negative in clownfish. (**Fig. 5B**)

Because our original PRDI metric is based on correlations calculated across many genomic windows, it does not provide high-resolution information about how much recombination evolved within any specific region of the genome. To address this, we developed a complementary measure, the local-PRDI (*PRDI_l_*). Briefly, for each recombination map, we ranked 2-kb genomic windows by recombination rates within each chromosome, and for each pairwise comparison, we quantified recombination landscape difference as the distance in ranks for each window. Mirroring the original PRDI approach, we then compared between-population and within-population comparisons and obtained an empirical value (*d_me_local_*; see Methods for details). We then subtracted from this empirical value a window-specific mean obtained from 20 neutral simulations (*d_ms_local_*), while the standard deviation across the simulations allowed us to assess statistical significance (**Fig. 5C**).

Given the general tendency of greater scaled PRDI in comparisons with higher inferred gene flow (**Fig. 4C-D**), we asked whether comparisons with higher inferred gene flow also show a stronger association between local recombination divergence (*PRDI_l_*) and local genetic differentiation. We quantified the association for each comparison as the Spearman rank correlation between *PRDI_l_* and F_ST_ (used here as a proxy for response to divergent selection) across 50-kb and 100-kb genomic regions genome-wide (see Methods for details; **SFig. 6**). Across comparisons, the association tends to increase with inferred gene flow at both scales of genomic resolution (50kb: Spearman *ρ* = 0.68; 100kb: Spearman *ρ* = 0.82; **Fig. 5D**). Although weaker, these trends remained positive after excluding the *A. calliptera* comparison (50kb: Spearman *ρ* = 0.49; 100kb: Spearman *ρ* = 0.71). We tested the statistical significance of this association using a linear mixed-effects model, with F_ST_, gene flow, and their interaction as fixed effects, and comparison (population or species pair) and chromosome as random effects. Consistent with the qualitative pattern observed in Figure 5D, the interaction term was positive (estimate=625, t= 3.510), suggesting that higher gene flow is associated with a more positive relationship between F_ST_ and *PRDI_l_*, and adding the interaction significantly improved the fit of the model compared with a model without interaction (likelihood ratio test: *χ*^2^ = 12.16, p < 0.001).

**Figure 6:**
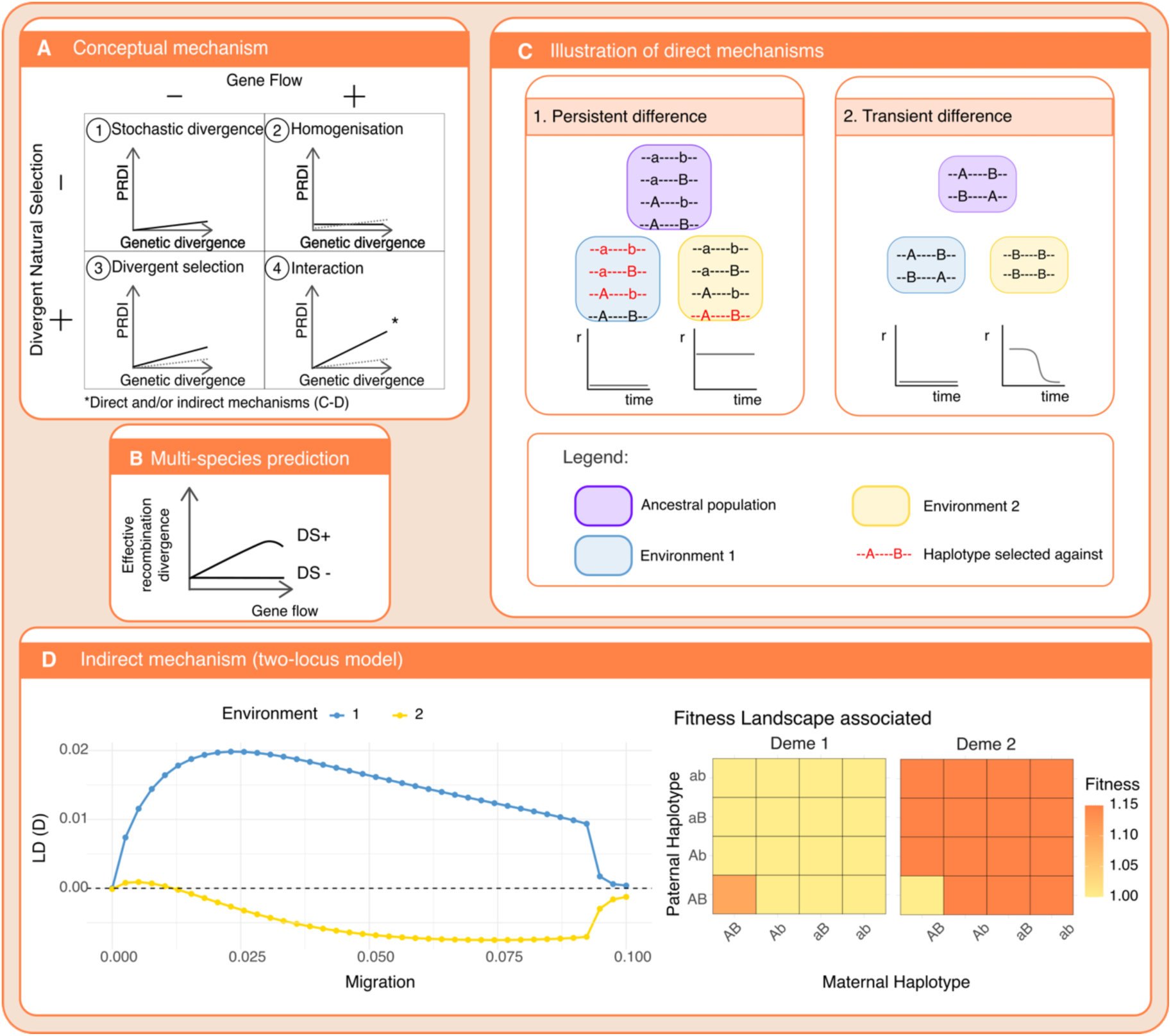
Concepts and mechanisms through which gene flow and divergent selection can generate divergence in effective recombination landscapes. **(A)** Qualitative expectations for population recombination landscape divergence (PRDI) across four evolutionary contexts. Rows indicate the absence or presence of divergent selection (DS), and columns the presence or absence of gene flow (GF). Grey dashed lines represent the null expectation (DS-/GF-). (1) In the absence of divergent selection and gene flow, effective recombination landscapes are expected to diverge through stochastic processes, (2) while gene flow tends to homogenise effective recombination landscapes between populations. (3) Divergent selection can alter LD differently between populations, thereby generating divergence. (4) Migration-selection dynamics enhance effective recombination landscape divergence by generating or maintaining environment-specific LD (panel D) or through direct changes in crossover rates and distribution (panel C). **(B)** Conceptual relationship between gene flow and effective recombination landscape divergence. Without divergent selection (DS-), gene flow is expected to homogenise effective recombination landscapes. With divergent selection (DS+), divergence increases, but at high levels of gene flow, migration overrules divergent selection, leading to a decrease in divergence. **(C)** Two examples of direct mechanisms involving changes in the underlying recombination rate r. Selection on recombination modifiers can produce either (1) persistent differences between environments when modifiers maintain pre-existing locally adapted haplotypes, or (2) transient differences when temporarily increasing recombination in one population facilitates the assembly of new advantageous haplotypes from standing variation. **(D)** Indirect mechanism illustrated using a two-locus, two-populations model in which the recombination rate is constant. Divergent selection and migration generate divergent LD between environments, thereby contributing to divergence in effective recombination landscapes. At higher migration (m=0.1), homogenisation causes LD to approach zero in both demes.

## Discussion

In this study, we quantified effective recombination landscape divergence across seven pairs of populations and species from four percomorph fish lineages, using the PRDI framework developed in Talbi et al. (2025). Divergence in the effective recombination landscape generally increased with genetic differentiation and divergence time. However, the amount of recombination landscape divergence per unit of genetic differentiation or divergence time varied substantially among comparisons (**Fig. 4A**, **SFig. 4A**, and **SFig. 5A**). Additionally, comparisons with higher inferred gene flow tended to show greater accumulation of PRDI per unit of genetic divergence or divergence time (**Fig. 4C**, **SFig. 4C**, and **SFig. 5C**). By developing a complementary local metric, the local PRDI (*PRDI_l_*), we revealed heterogeneity of effective recombination divergence along the genome and identified rapidly (and slowly) evolving regions (**Fig. 5**). Although based on a limited number of independent comparisons (n = 7), the consistency of these patterns across lineages suggests that genetic differentiation, divergence time, and gene flow may be important to understand the evolution of recombination landscape divergence.

We emphasize that both PRDI and *PRDI_l_* rely on LD-based recombination maps and therefore reflect the effective recombination landscape, not the physical crossover landscape alone, as would be observed in the gametes of present-day individuals (Peñalba and Wolf 2020). LD-based estimates of *ρ* integrate the combined effects of crossover rates, selection on linked loci, and *N_e_*, and, although we explicitly model demography, we do not fully separate changes in crossover frequency from changes in genealogical structure driven by selection and migration. However, from an evolutionary perspective, it is this effective recombination landscape – emerging from the interaction of crossovers, selection, and migration – that is of great interest, as it governs how alleles interact across the genome and how locally adapted haplotypes persist or break down.

PRDI was positive across all seven comparisons, including the youngest species and population pairs which diverged within approximately 20,000 years: the Victoria and Masoko cichlids, and the stickleback pairs. This indicates that detectable divergence in effective recombination landscapes can arise over relatively short evolutionary timescales in percomorph fishes. This pattern does not necessarily imply equally rapid evolution of the underlying crossover landscapes. Nevertheless, it is notable that effective recombination landscapes differ among closely related populations despite the absence of a functional PRDM9-directed recombination mechanism in these lineages (Singhal et al. 2015; Baker et al. 2017). Differences in both crossover distribution and population recombination landscape have been previously suggested between marine and freshwater stickleback based on pedigrees (Venu et al. 2024) and on low hotspot overlap inferred from LD-based maps (Shanfelter et al. 2019), while in Talbi et al. (2025), we examined the divergence in recombination landscapes in the Lake Masoko cichlid pair. Our results confirm and extend this picture across percomorph fishes while explicitly accounting for variation arising from genetic drift and demography.

Our local-PRDI analysis further shows substantial heterogeneity in recombination landscape divergence along the genome. Some genomic regions display significantly stronger divergence than expected under neutral evolution, whereas other regions remain relatively conserved with respect to their recombination landscape (**Fig. 5C**). This heterogeneity in recombination divergence along the genome suggests that multiple mechanisms may contribute, including local genomic features, larger structural variants, and broader differences in genomic architecture. Comparative work across fishes has, for example, implicated karyotype variation in heterochiasmy and identified associations between recombination rates and CpG content (Kivioja and Rastas 2024). Combining local-PRDI with molecular and structural genomic analyses may therefore help identify the genomic features and mechanisms that shape recombination landscape divergence.

A central result of our study is that effective recombination landscape divergence accumulates at markedly different rates per unit of genetic differentiation or divergence time across different population and species pairs. Pairs with relatively lower genetic differentiation tend to have a greater rate of PRDI accumulation per unit of genetic distance, which suggests that effective recombination landscape divergence may not accumulate linearly with genetic differentiation (**Fig. 4**, **Fig. S4**) or through time (**Fig. S5**), but instead may exhibit more complex evolutionary dynamics. Although our dataset includes only seven independent comparisons, limiting statistical power, the amount of recombination landscape divergence accumulated per unit of genetic divergence or divergence time tended to be higher in pairs with greater gene flow. Additionally, the local analyses showed a related pattern: the association between local recombination divergence and local genetic differentiation tended to be stronger in comparisons with greater inferred gene flow. Together, these observations suggest that gene flow may contribute to heterogeneity in effective recombination-landscape divergence among systems.

This relationship is not an automatic consequence of migration. In the absence of divergent selection, gene flow would generally be expected to homogenise populations, including their LD structure and effective recombination landscapes, leading to low and stable PRDI values (Samuk and Noor 2022). Conversely, in the absence of gene flow, divergent selection could increase PRDI locally when the genomic regions targeted by selection differ between populations. When gene flow occurs between populations experiencing divergent selection, however, migration continually introduces ancestry blocks into genetic backgrounds in which their fitness effects differ. The resulting migration–selection dynamics can generate or maintain population-specific LD patterns and thereby increase divergence in effective recombination landscapes (**Fig. 6A**).

Together, these scenarios suggest that the effect of gene flow on effective recombination landscape divergence depends on the presence of divergent selection (**Fig. 6 B**). Such patterns may arise through direct changes in crossover distribution or genomic architecture, including the evolution or fixation of recombination modifiers, inversions, and other structural variants (**Fig. 6C**; Kirkpatrick and Barton 2006; Ortiz-Barrientos et al. 2016) or indirectly through changes in population-specific LD (**Fig. 6D**).

Our two-locus model (**Fig. 6D**) illustrates the indirect mechanism – i.e., a mechanism that impacts the LD structure without changing the underlying crossover distribution. Despite a constant crossover rate, divergent selection and migration generate LD in opposite directions in the two environments, with the strongest divergence arising at intermediate levels of migration. Effective recombination landscapes can therefore diverge even without changes in the underlying crossover distribution. This result provides a simple interpretation of the empirical tendency for the association between local-PRDI and local genetic differentiation to become stronger with increasing gene flow.

Several limitations of this study should be considered. First, statistical power is limited due to the small number of independent comparisons, and the strength of some relationships is influenced by one or a few data points. Second, inferred migration rates depend on the demographic models used, although it is worth keeping in mind that our interpretation relies on relative differences among systems rather than absolute estimates. Third, while the broad taxonomic scope of the study highlights the consistency of several patterns across evolutionarily disparate taxa – suggesting that they may reflect general features of recombination landscape evolution – it also makes it difficult to disentangle individual causal factors underlying these patterns. Future studies based on replicated contrasts among closely related populations will therefore be important for isolating the contributions of specific evolutionary processes. In particular, comparisons between sympatric and allopatric population pairs, which differ in the presence or absence of gene flow, as well as populations evolving in similar versus contrasting environments, will provide a powerful framework for identifying specific underlying causes. Finally, integrating LD-based maps with pedigree- or gamete-based recombination maps would help distinguish how much of the divergence reflects changes in crossover rates versus selection-driven changes in genealogical structure. Initial steps in this direction have been taken by (Delaive et al. 2026).

Overall, our results demonstrate that effective recombination landscapes are not static genomic backdrops but dynamic features that diverge among closely related populations and species, and that this divergence varies substantially among genomic regions and may be shaped by the interaction between divergent selection and gene flow. By quantifying divergence at both genome-wide and local scales, PRDI provides a framework for incorporating effective recombination-landscape evolution into studies of adaptation and speciation. We further envisage that this work will motivate theoretical studies on the evolution of effective recombination landscapes.

## Materials and Methods

### Sampling, DNA extraction, and sequencing

For this project, we generated data for a total of 637 individuals from 14 populations across 8 different species. We extracted DNA from 43 *Pundamilia nyererei* individuals sampled in Lake Victoria, in Tanzania, 36 *Astatotilapia nubila* individuals sampled in a swamp near Lake Victoria, and 83 *Nothobranchius furzeri* individuals sampled in two different river basins in Mozambique, namely Chefu and Limpopo. We used the PureLink™ Genomic DNA Mini Kit, following the standard protocol. The extracted DNA was then sequenced on the Illumina NovaSeq 6000 platform, obtaining 150bp paired-end reads.

We also sequenced paired-end read data from DNA already extracted for 39 and 37 individuals from two species of Lake Tanganyika cichlids, respectively *Jabarichromis pfefferi* and *Pseudosimochromis curvifrons*. Finally, we also obtained paired-end read data already sequenced for 48 and 38 individuals from Papua New Guinea and Indonesia (specifically from Sulawesi) for *Amphiprion clarkii*. We also included 70 and 69 individuals from two ecotype populations, benthic and littoral, from the cichlid species *Astatotilapia calliptera* from Lake Masoko, from previous work (Talbi et al. 2025). Similarly, we included 35, 48, 44, and 47 individuals from Finger Lake, Long Lake, Spirit Lake, and Watson Lake populations of Alaskan threespine sticklebacks (*Gasterosteus aculeatus*) that were used in previous work (Hendry et al. 2024; Malinsky et al. 2025).

### Read alignment, variant calling, and filtering

The short-read data from each of the four taxa groups (cichlids, stickleback, killifish, and clownfish) were mapped to a separate reference genome. All cichlids were aligned to the *Astatotilapia calliptera* reference genome (GenBank ID: GCA_900246225.6), which enabled us to merge the VCFs and facilitate comparisons across the group. Specifically, the Victoria cichlids *Astatotilapia nubila* and *Pundamilia nyererei* were aligned jointly to the reference genome, while the Tanganyika cichlids *Jabarichromis pfefferi* and *Pseudosimochromis curvifrons* were aligned independently to the reference genome. *Amphiprion clarkii* populations were aligned to the *Amphiprion clarkii* reference genome (GenBank ID: GCA_027123335.1), and *Nothobranchius furzeri* populations were aligned to the *Nothobranchius furzeri* reference genome (GenBank ID: GCA_001465895.2). Finally, for the threespine sticklebacks, we used the *Gasterosteus aculeatus* reference genome v.5 (GenBank ID: GCA_016920845.1) (**Supplementary Table 1**).

We aligned reads and called variants for five different sets of populations: (i) *Amphiprion clarkii* from Indonesia and Papua New Guinea population, (ii) *Nothobranchius furzeri* from Chefu and Limpopo, (iii) *Pundamilia nyererei* and *Astatotilapia nubila*, (iv) *Jabarichromis pfefferi,* and (v) *Pseudosimochromis curvifrons* from Lake Tanganyika. As mentioned above, we also included individuals from published work for *Astatotilapia calliptera* from Lake Masoko, and for four populations of *Gasterosteus aculeatus* from Alaska. For these two last sets of populations, we used the VCFs that we had already generated. The variant calling and filtering methods were the same as described below.

To align reads for the five sets of populations listed above, we used bwa-mem v0.7.17. (Li 2013). We then used the MarkDuplicates tool from the Picard package v.2.26.6 to tag PCR and optical duplicate reads. Variant calling was performed with GATK v.4.2.3 (DePristo et al. 2011), using the HaplotypeCaller in GVCF mode for each individual separately. We then performed joint genotyping of all samples within each set using GenotypeGVCFs with the --include-non-variant-sites option. These ‘all sites’ VCF files were used to generate callability by filtering out: (i) sites outside the specific depth range (**Supplementary Table 1)**, (ii) sites where more than 10% individuals have missing data, (iii) sites flagged as low quality by GATK and (iv) sites with poor mappability (see Talbi et al., 2025). The proportion of masked sites represented between 32.8 and 45.2% of the genome (**Supplementary Table 1**). Finally, we applied hard filters following GATK best practices. These included filtering out variants with mapping quality (MQ < 40), strand bias (FS > 40), quality normalized by depth (QD) < 2, and excess heterozygosity as compared to Hardy–Weinberg expectations (ExcessHet > 40). Finally, we used vcftools to remove indels and keep only biallelic sites with the --remove-indels, and --min-alleles 2 --max-alleles options respectively.

To obtain a VCF with all cichlid populations, we merged the four VCFs (i.e., (i) *A. calliptera* from Masoko, (ii) *P. nyererei* and *A. nubila* from Victoria, (iii) *J. pfefferi*, and (iv) *P. curvifrons* from Tanganyika) that have been aligned to the *A. calliptera* reference genome using the `merge` command from bcftools v.1.16, with the `-missing-to-ref` option. Then, we created a joint mask by combining the masks from all five species, excluding all non-callable sites from the final VCF. This final VCF resulted in 282 individuals and 31 536 364 SNPs.

As a result of the above procedure, we ended up with four VCF files (datasets: the cichlids, the sticklebacks, the killifish, and the clownfish) with biallelic SNPs. These datasets served as the input for all the following analyses. Summary statistics for these datasets are provided in **Supplementary Table 1**.

### NJ trees and sample selection

For each dataset, we constructed a neighbor-joining tree based on a genetic distance matrix, calculated as the average number of single-nucleotide differences among all individuals. Pairwise difference matrices were divided by the size of the callable genome to obtain pairwise distances per base pair, which were then used as input for the nj() tree-building function implemented in the package *ape* in R (Paradis et al., 2004). Individuals who were misassigned or showed excessive relatedness to others were excluded from the analyses. Specifically, we excluded 8 individuals from *P. curvifrons*, one from *J. pfefferi*, one from *P. nyererei*, one from the Indonesian population of *A. clarkii*, and none from the Papua New Guinean population of *A. clarkii*, *A. nubila*, any populations of *G. aculeatus*, *or N. furzeri*. From the final list of individuals, we created for each population a VCF using the `–view` command from bcftools v.1.16.

### PCA

To obtain the genetic structure for each of the datasets, we performed principal component analysis (PCA) using smartPCA (Patterson et al. 2006) on data filtered for minor allele frequency >=0.05 with plink v1.9 (Purcell et al. 2007) using the --maf 0.05 option. Linkage disequilibrium (LD) pruning was applied using the bcftools plugin +prune (Danecek et al. 2021) with the -m 0.8 -w 1000 options.

### Inference of demographic history and estimation of the level of gene flow

For all demographic analyses, we aimed to reduce the effects of linked selection by using only sites from noncoding regions of the genome, masking annotated exons, introns, and promoters. For each dataset, we used the available reference genome annotations, which are listed in the **Supplementary Table 1**. To estimate split times between population pairs and past changes in effective population size (*N_e_*), we used smc++ v.1.15.4 (Terhorst et al. 2017), using the following commands: vcf2smc, estimate, split, and plot. Smc++ scales inferred effective population size and coalescent times conditionally on the specified mutation rate. Therefore, we provided smc++ with taxon-specific mutation rates (*μ*) obtained from the literature, with the corresponding values and references listed in **Supplementary Table 1.** The same mutation rates were used to convert the time axis into generations.

We used easySFS, a Python utility built on the dadi libraries (Gutenkunst et al. 2010) to obtain the joint minor allele frequency spectrum (joint MAF). We then used fastsimcoal2.7 (Excoffier et al. 2021) to estimate the level of gene flow for each pair of populations. The split time and changes in *N_e_* inferred by smc++ were used as fixed parameters. The input file for fsc2 was converted directly from the smc++ output using a custom Python script. We estimated asymmetrical migration rates following the split using a logarithmic scale before allowing the program to explore a broader range of migration-rate values. This reduces the chance that the analysis stays trapped in a local likelihood optimum. We ran 30 simulations for each population pair, each with different starting parameters to explore different regions of parameter space. For each population pair, we retained the directional migration-rate estimates from the run with the highest likelihood among the 30 runs. We then reported the migration rate for each population pair as the average of the two directional gene flow estimates.

### Subsampling for recombination analyses

We used the `shuf-n` command to randomly draw a set of n individuals for the first subset and the command `grep-v` to obtain a second subset b with the *N* − *n* remaining individuals for each population. These subsets a and b are composed of non-overlapping sets of individuals and act as biological replicates (see Talbi et al., 2025). To choose n, we divided N by two and rounded the number above. For each subset, we generated a separate VCF using the bcftools v.1.16 `view` command and used these VCFs for recombination rate inference.

### Inference of recombination rates

We used the software *pyrho* to infer recombination rates along the genome from patterns of Linkage disequilibrium (LD) present in the data, while accounting for demography (Spence & Song, 2019). First, we used the `make_table` command to build likelihood tables between each pair of biallelic sites, incorporating the mutation rate mu (*μ*) and demographic changes inferred by smc++ for each population subset. We used the `–approx` `flag` with `–moran_pop_size` N, where N was set to 1.5 times the number of haplotypes in each subset.

Next, we ran the –`optimize` command in *pyrho* on each VCF file using the previously computed likelihood tables. To identify the optimal value for the block penalty parameter, we relied on previous simulations based on different evolutionary scenarios (see Talbi et al., 2025). As the different subsets had a similar number of individuals (∼ 20 individuals) and comparable mutation rates, we selected a block penalty value of 15, which minimized both false positives and false negatives in the simulations. This value was used in all runs with a window size of 50 SNPs. For each population subset, *pyrho* produced estimates of recombination rates between each pair of SNPs.

### Neutral coalescent simulations

To obtain a neutral baseline for the differences in recombination expected under the effect of the demography alone, we applied the same approach as in Talbi et al. (2025) to all seven population pairs. Briefly, we used msprime v.1.0.2 (Baumdicker et al., 2021) to simulate genetic data, using the demographic history parameters inferred for the population pair from the empirical data with smc++ and using a unique underlying recombination map within each simulation. This map corresponds to the inferred recombination map of one of the two populations or species pair. The Python script used to run these simulations is available from GitHub (https://github.com/MarionTalbi/RecombinationLandscapesAcrossPercomorphFishes). For each population pair, we ran one simulation per chromosome (between 19 and 24 regarding the species) and sampled the same number of individuals as in the empirical data to output one final VCF file. We then randomly subsampled the (a) and (b) subsets and processed the output similarly to what we did for the empirical data.

### Correlation in recombination rates and PRDI calculation

We obtained the mean recombination rates in 2 kb windows for all datasets using the `PhysicalWindowAverages` command from the evo package v.0.1 r28. We processed each dataset separately (i.e., cichlids, sticklebacks, clownfish, and killifish) and applied the same analysis to both the empirical and simulated data. The correlations and map distances were then calculated using R scripts, available on GitHub (https://github.com/MarionTalbi/RecombinationLandscapesAcrossPercomorphFishes).

To quantify the amount of recombination rate evolution between each population pair, we used the Population Recombination Divergence Index (PRDI). Briefly, this measure is obtained by comparing the differences between the median of recombination rates correlation between- and within-populations, calculated from empirical (d_me_) and simulated (d_ms_) data (See Talbi et al., 2025 for more details and Figure 1). We then calculated PRDI as the difference between these value (i.e., dm_e_ - dm_s_) at the genome and chromosome levels and performed the statistical analyses using a custom R script. Scaled PRDI values were obtained by dividing genome-wide PRDI by estimates of F_ST_, d_a_, and divergence time.

To assess whether PRDI was significantly different from zero, we used a bootstrap method. More precisely, for each comparison, we first calculated the six within- and between-population recombination map correlations across 500 kb genomic windows, either genome-wide or separately for each chromosome, as needed to calculate PRDI. We then generated bootstrap datasets by resampling these genomic windows with replacement, keeping the same number of windows as in the original dataset. We then recalculated dm_e_ and dm_s_ from the resampled correlations and calculated PRDI as dm_e_ - dm_s_. We repeated this procedure 1000 times, and the 95% bootstrap confidence interval was defined by the 2.5th and the 97.5th percentiles of the resulting distribution. PRDI was considered significantly different from zero when the interval didn’t include zero.

We also quantified PRDI heterogeneity along the genome. To do so, we repeated the neutral simulation procedure described above 20 times for each of the seven population pairs and calculated the mean recombination rates in non-overlapping 2 kb windows. Repeating the simulation allowed us to average over stochastic variation in ancestry and to obtain a standard deviation for each window.

For each dataset, we then worked chromosome by chromosome and ranked all the 2-kb windows according to their recombination rate. We used ranking rather than absolute recombination values to capture relative changes along the genome without being influenced by differences in the absolute recombination rate.

Analogous to the calculation of the original PRDI, we computed the absolute differences in rank within each population, denoted Δ*r*_*intra*1_ and Δ*r*_*intra*2_, and between the populations, yielding four values, denoted Δ*r*_*inter*1,1_, Δ*r*_*inter*1,2_, Δ*r*_*inter*2,1_, Δ*r*_*inter*2,2_ We then took the minimum of the two intra-population and averaged the four inter-population values and defined:

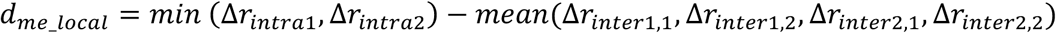

This value *d_me_local_* is obtained from the empirical data. We applied the same procedure to each of the 20 simulation replicates to obtain *d_me_local_*, for each simulation *s*. Then, we calculated the mean and the standard deviation across simulations.

Finally, we calculate the local-PRDI (*PRDI_l_*), for each 2 kb window as:

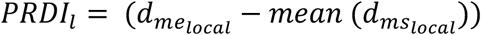

We then derived a lower and an upper bound to quantify the expected neutral range, defined as:

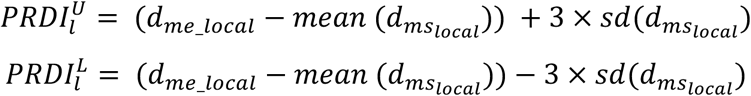

### Measures of genetic differentiation

To quantify the level of genetic differentiation between each population pair, we calculated F_ST_, based on allele frequency differentiation, d_a_, the net nucleotide difference between two populations, calculated as 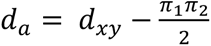 (Cruickshank and Hahn 2014), and *π*, the nucleotide diversity calculated within each population. We used the `F`_ST_ command from the evogenSuite software to compute these statistics. For the nucleotide diversity, we considered the number of callable sites with the `–` `AccessibleGenomeBD` option.

### Genome-wide association between selection and recombination divergence

For each pairwise comparison, we calculated *l_PRDI_* and F_ST_ in 2 kb windows along the genome. Then, we aggregated these estimates by averaging across 2 kb windows within larger 50 kb and 100 kb non-overlapping windows for both values, producing 50 kb and 100 kb binned values for both metrics.

To quantify the strength of the association between *l_PRDI_* and F_ST_ within each comparison, we first calculated Spearman rank correlation coefficients because both variables deviated from normality according to Shapiro-Wilk tests (p-value<0.05). We then tested whether these correlation coefficients were associated with inferred gene flow across comparisons, again using Spearman rank correlation. This p-value was not used because it does not incorporate uncertainty associated with the within-comparison correlation estimates. Therefore, we additionally used linear-mixed effects models to formally test whether gene flow affected the relationship between *l_PRDI_* and F_ST_.

Linear mixed-effects models were fitted using the lmer() function from the lme4 package in R. For each window size, we compared two models:

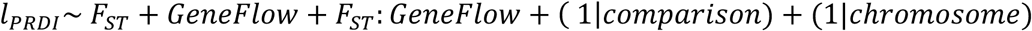

and

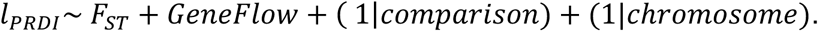

The first model included an interaction between F_ST_ and gene flow, and the second model excluded this interaction. Both models included comparison and chromosome as random effects. We compared models fit using a likelihood ratio test with the anova() function, to test whether including the interaction term improved the model.

### Two-Locus Migration-Selection Equilibrium

To investigate how equilibrium linkage disequilibrium (LD) varies with migration rate, we used a custom R script to simulate the frequency dynamics of four haplotypes (AB, Ab, aB, and ab) in two populations connected by migration. The model followed a life-cycle of selection, recombination, and migration: selection acts on adults, recombination occurs during gamete formation, and migration takes place at the juvenile stage before individuals become adults in the next generation. The model included a deme-specific fitness matrix specifying the fitness of each diploid genotype and represented a scenario of divergent selection between demes. The model then reported the equilibrium LD obtained under different levels of gene flow. The R script is available on GitHub.

## Data and code availability

The inferred recombination maps from simulated and empirical data, as well as estimates of F_ST_, d_XY,_ and genetic diversity (*π*) are available on DataDryad (DOI: 10.5061/dryad.0gb5mkmhh). The raw sequence data are available under the following accessions: BioProject PRJNA1135142 (*Nothobranchius furzeri*), BioProject PRJEB25984 and the Tree of Life project <data_to_be_released> (*Jabarichromispfefferi* and *Pseudosimochromis curvifrons*), BioProject PRJEB27804 (*Astatotilapia calliptera* from Lake Masoko), BioProject PRJEB49185 (*Gasterosteus aculeatus*), and BioProject PRJNA1202426 (*Amphiprion clarkii*).

The R code used to analyse the recombination maps and to generate the two-locus model is available on GitHub (https://github.com/MarionTalbi/RecombinationLandscapesAcrossPercomorphFishes).

## Acknowledgements

We would like to thank members of Ole Seehausen’s group, particularly Pooja Singh, as well as members of Nicolas Salamin’s group, namely Anna Marcionnetti, Sarah Schmid, Lucy Fitzgerald and Alberto García Jimenez, for helpful discussions.

