## Supplementary materials for "Heterogeneous evolution of effective recombination landscapes across percomorph fishes"

**Affiliations:**

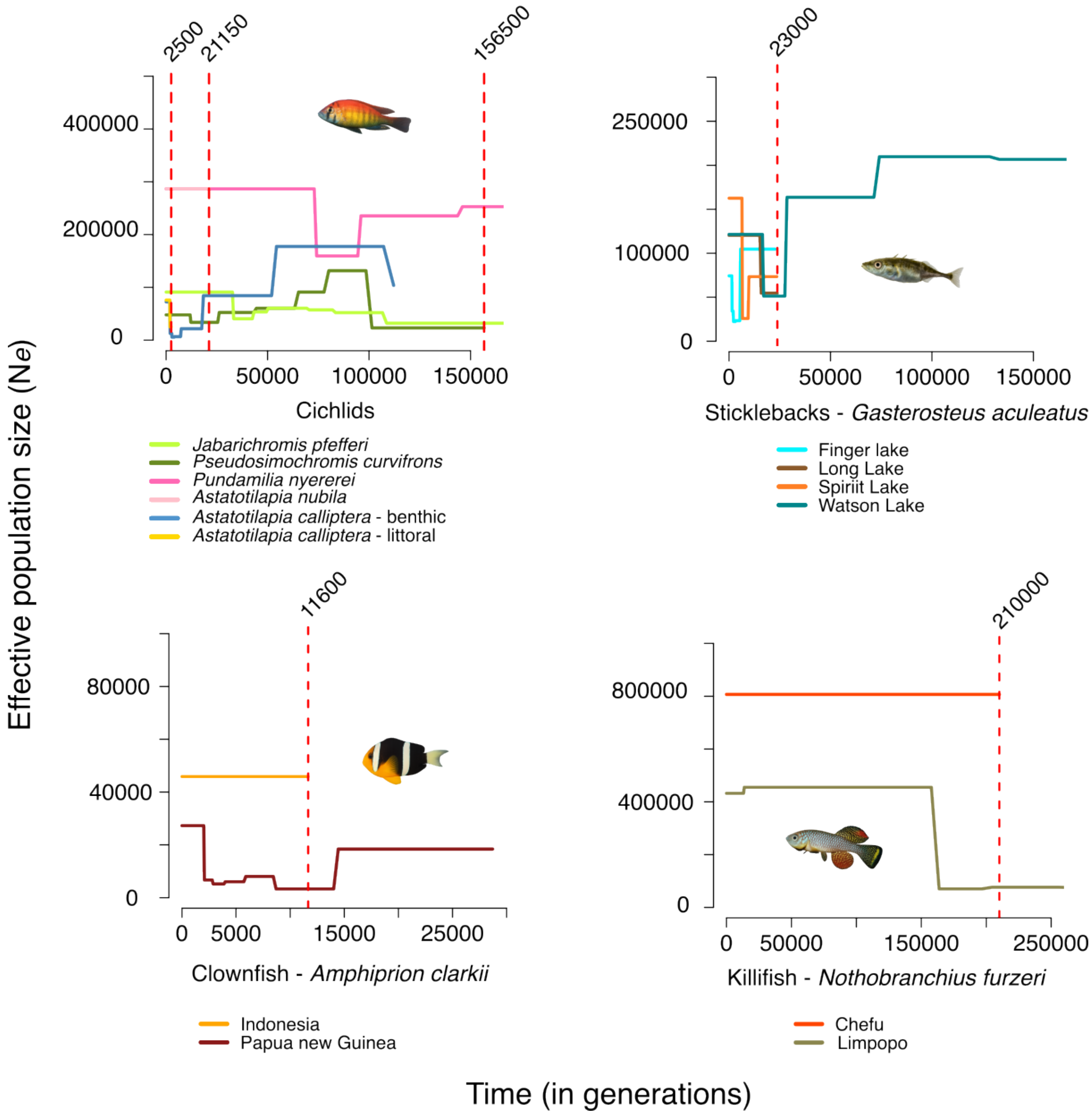


**Figure S1: Demographic history for each lineage**. Demographic history as inferred from smc++. This method relies on a Hidden Markov Model to estimate the time to the most recent common ancestor (TMRCA) conditional on the site frequency spectrum distribution *(Terhorst et al. 2017)* . The red vertical dashed lines represent the split time inferred with the same software, between each population pair. For the sticklebacks, it corresponds to the mean between the two inferred split times (**Table 1**).

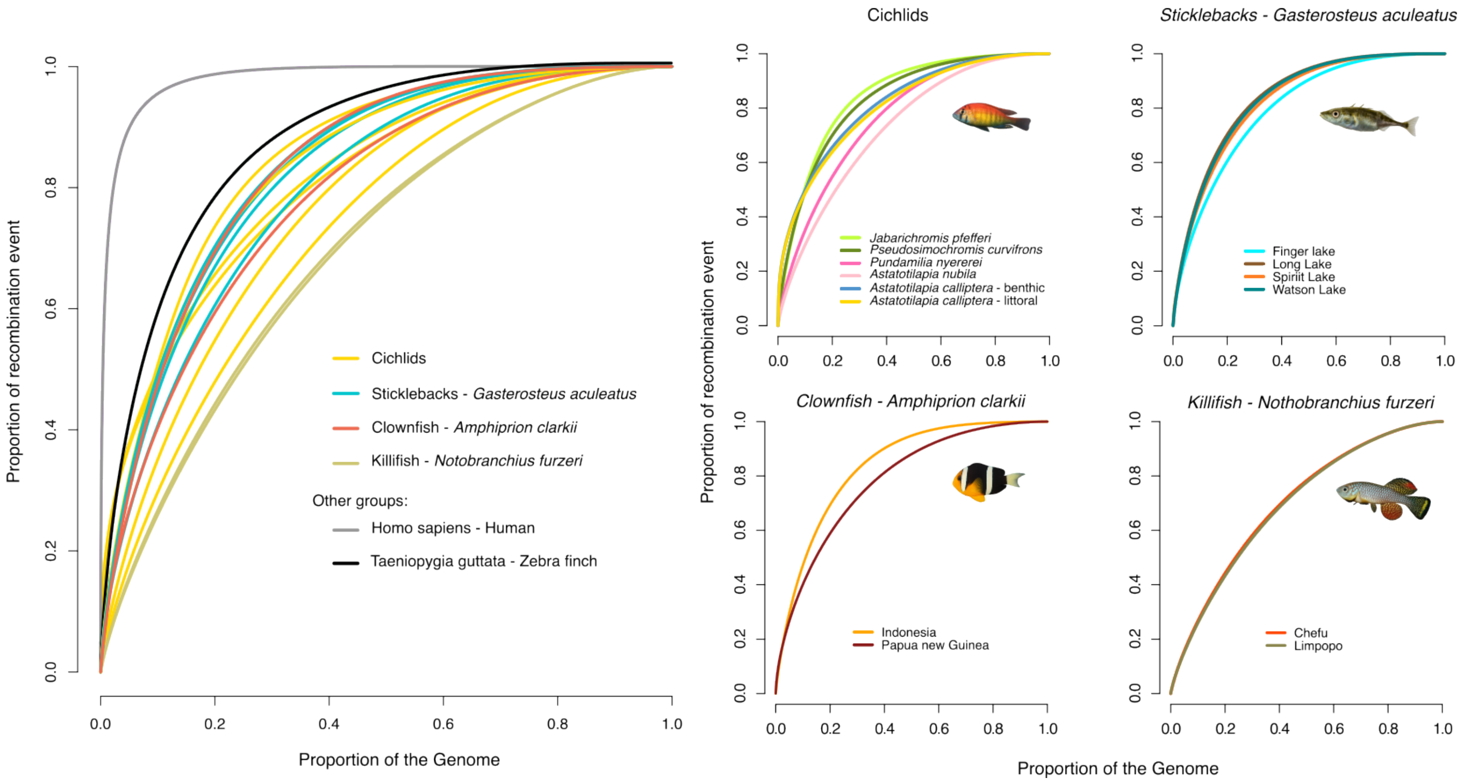


**Figure S2: Cumulative curve for each lineage independently.** Similarly to Fig. 4A, cumulative curves represent the number of recombination events per genome proportion for all 14 populations, including *Homo sapiens* (gray) and *Taeniopygia guttata* (black). Each of the four lineages is also shown separately at the right of the figure.

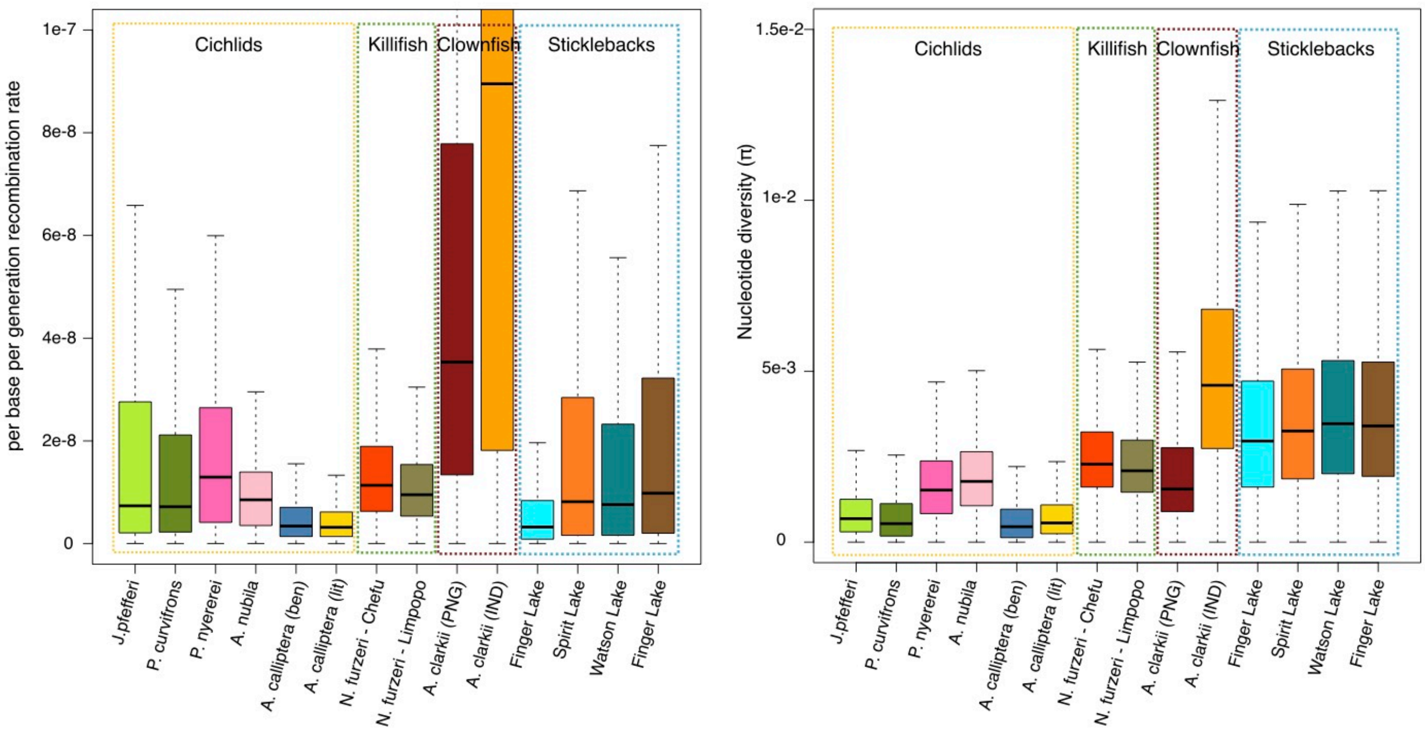


**Figure S3: Genome-wide effective recombination rate and nucleotide diversity.** For each population/species of this study, boxplots represents values for each 2kb window across the genome for (**left**) the per-base per-generation recombination rate and (**right**) the nucleotide diversity ($\pi$). Dashed rectangles separate the four focal groups.

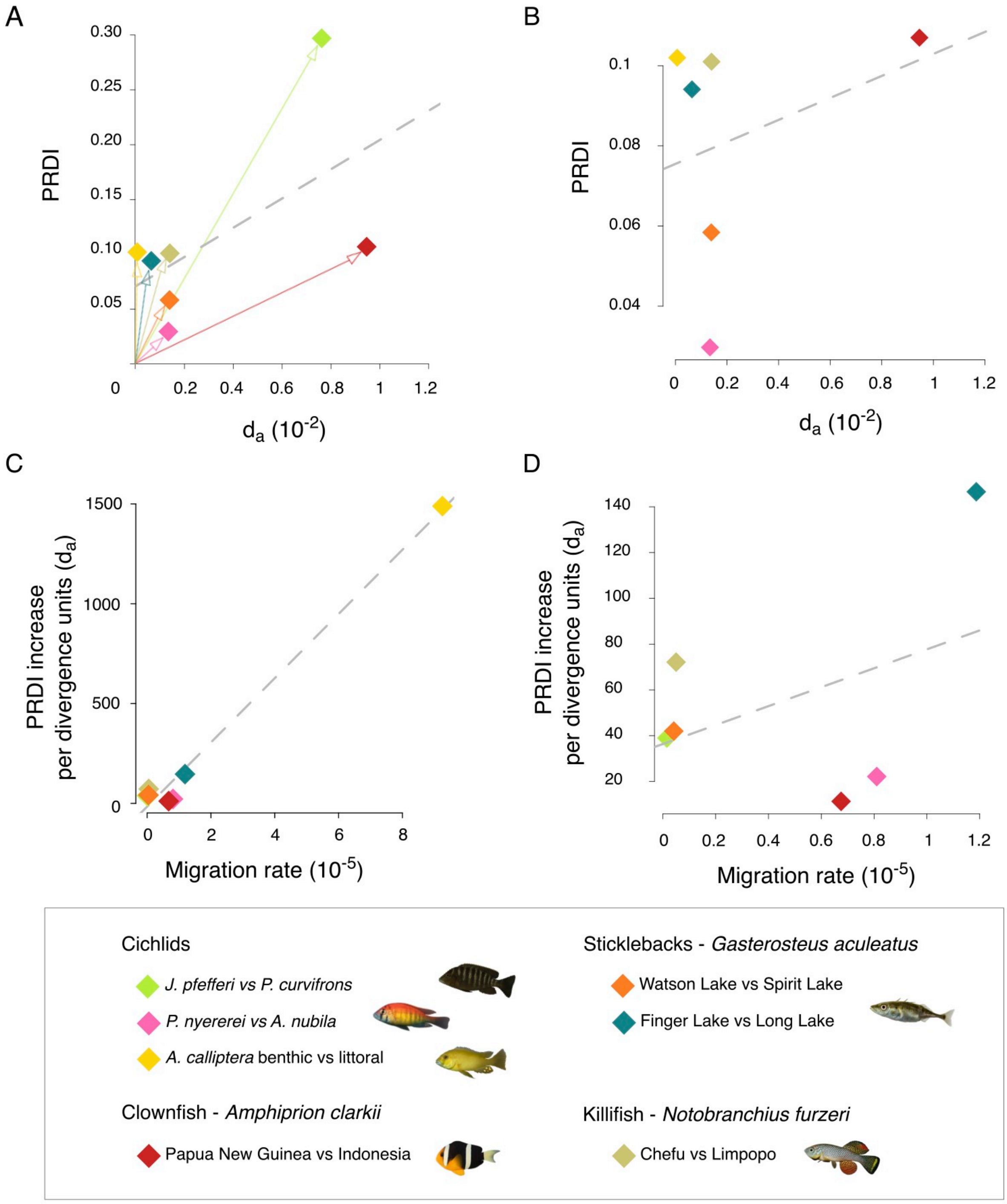


**Figure S4: Recombination landscape divergence and net genetic divergence. (A)** Relationship between the population recombination divergence index (PRDI) and mean genome-wide *net* genetic *divergence* ($d_a$) for each population pair. Panel (**B**) shows the same relationship without the *J. pfefferi* and *P. curvifrons* comparisons. Each coloured symbol represents one pair. Arrows from the origin highlight that the slope, representing PRDI per unit of divergence, differs among comparisons. **(C)** Relationship between the scaled recombination landscape divergence, calculated as PRDI divided by $d_a$, and migration rate. (**D**) Same relationship without the *A. calliptera* comparison.

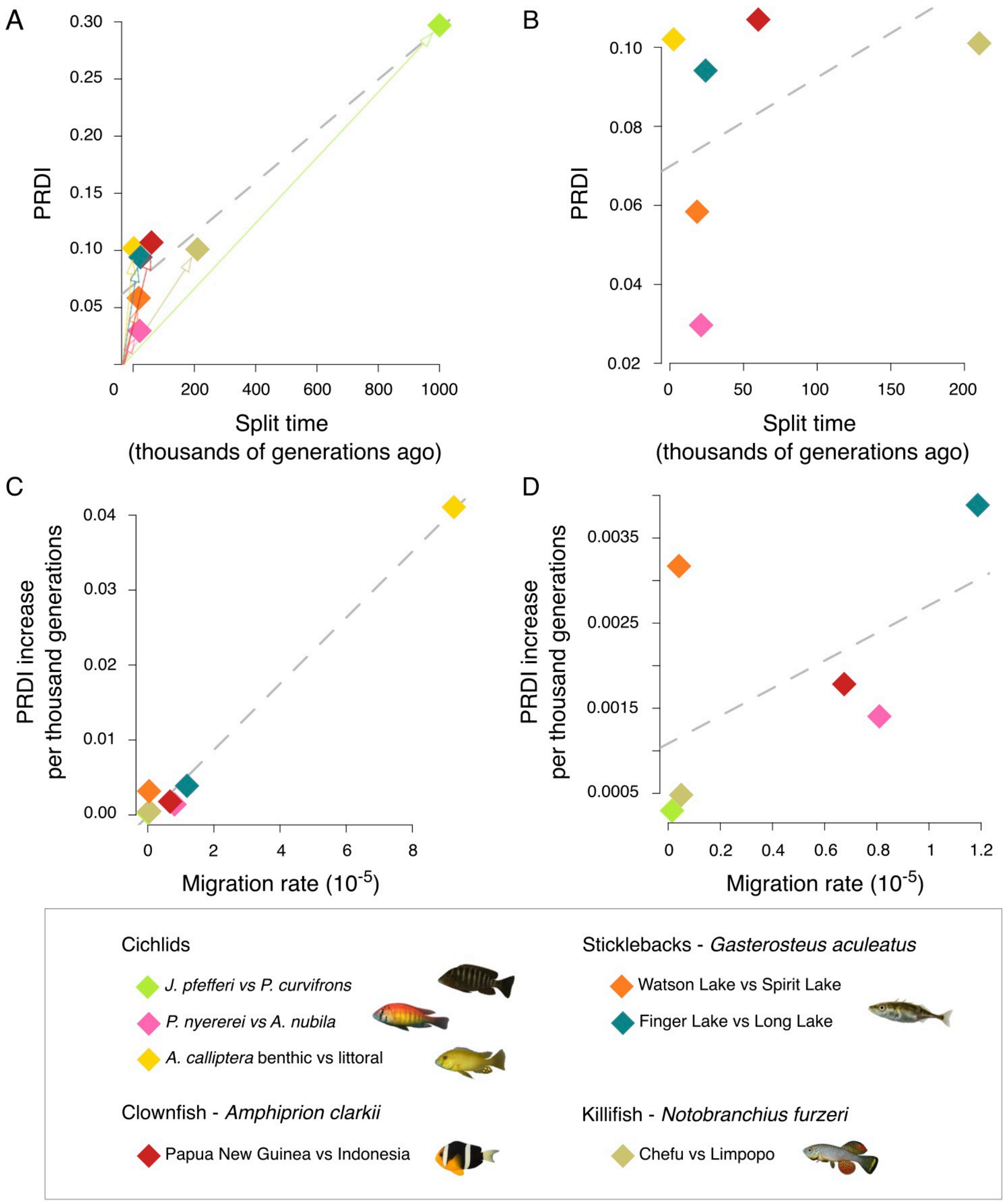


**Figure S5: Recombination landscape divergence and time since divergence from the common ancestor. (A)** Relationship between the population recombination divergence index (PRDI) and split time, in thousands of generations, inferred with smc++ and adjusted accordingly to current literature for the *J. pfefferi* vs *P. curvifrons* and the *A. clarkii* comparisons. This figure is organised similarly to Fig. 4 and SFig. 4, but focuses on split time from the common ancestor.

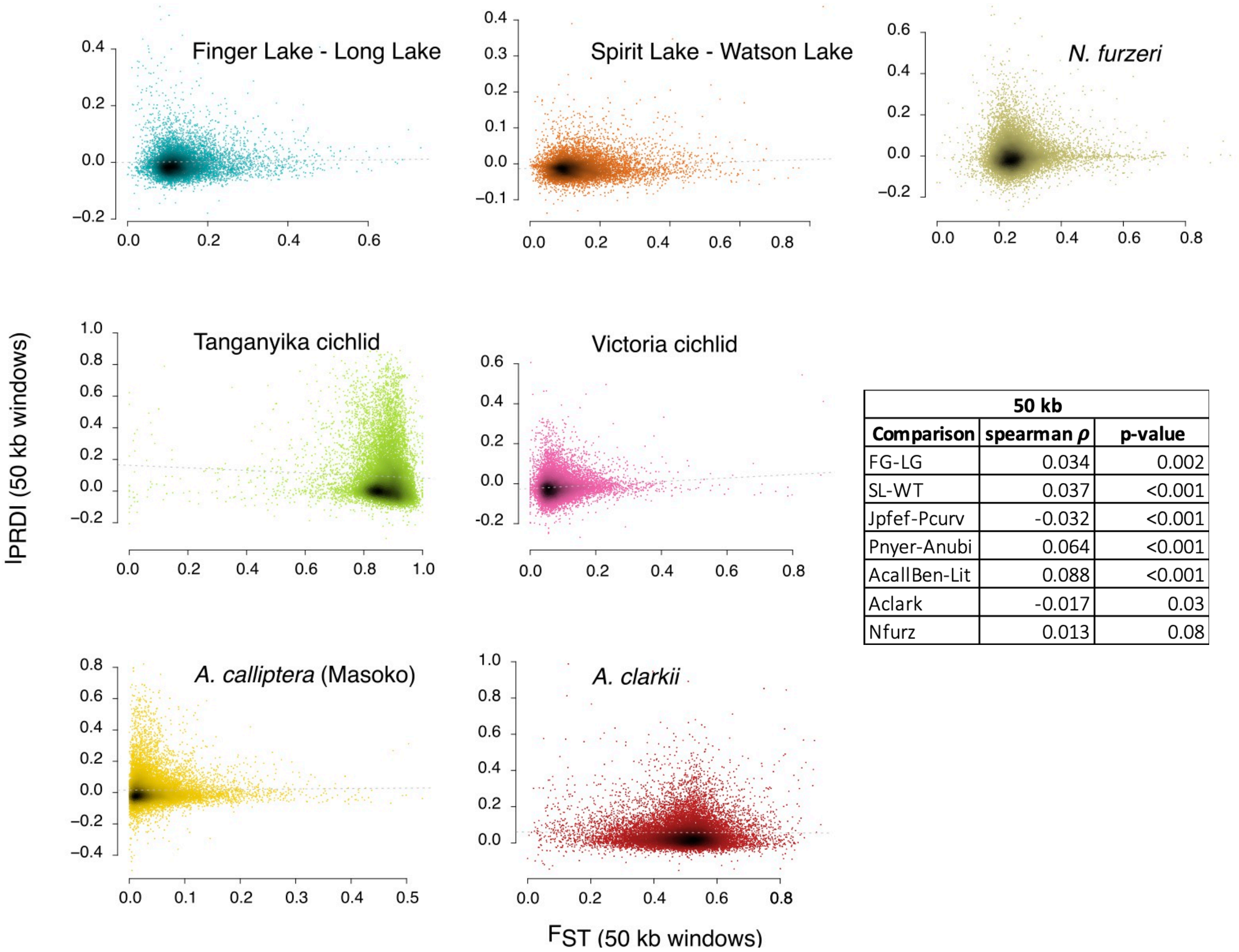


| 50 kb | | |
|---|---|---|
| **Comparison** | **spearman $\rho$** | **p-value** |
| FG-LG | 0.034 | 0.002 |
| SL-WT | 0.037 | <0.001 |
| Jpfef-Pcurv | -0.032 | <0.001 |
| Pnyer-Anubi | 0.064 | <0.001 |
| AcallBen-Lit | 0.088 | <0.001 |
| Aclark | -0.017 | 0.03 |
| Nfurz | 0.013 | 0.08 |

**Figure S6: Relationship between local recombination divergence ($PRDI_l$) and genetic differentiation ($F_{ST}$).** For each comparison, we computed Spearman rank correlation between $PRDI_l$ and $F_{ST}$ across 50 kb genomic windows genome-wide. Window-level $PRDI_l$ and $F_{ST}$ were obtained by averaging 2kb estimates within each 50 kb window. Spearman $\rho$ and its associated p-value are reported in the table.

**Table S1: Information on each dataset used in this study.** This table refers to the different species and populations present in each dataset. We also listed the accession number for the reference genomes used, the mutation rate used in the analyses, the depth that we used for filtering, as well as the final number of individuals per VCF and the final number of single nucleotide polymorphisms (SNPs). Mutation rates were obtained from (Marques et al. 2025) for the Victoria cichlids, (Malinsky et al. 2018) for Masoko and Tanganyika cichlids, (Cui et al. 2019) for killifish, (Delrieu-Trottin et al. 2017) for clownfish, and (Zhang et al. 2024) for sticklebacks.

| Dataset | Comparison group | Reference genome | Reference Genome Accession Number | Genome annotation Accession Number | SNPs number | Genome size (Mb) | Mutation rate | Population / Species | Depth filter (min - max) | Final number of individuals |
|---|---|---|---|---|---|---|---|---|---|---|
| Cichlids | Victoria's cichlid | *Astatotilapia calliptera* | GCA_900246225.6 | GCF_900246225.1 (lift over) | 31536364 | 880 | 2.4E-09 | *Pundamilia nyererei* | (700 - 27th ; 1600 - 99th) | 41 |
| | | | | | | | | *Astatotilapia nubila* | (700 - 27th ; 1600 - 99th) | 36 |
| | Tanganyika cichlids | | | | | | 3.5E-09 | *Jabarichromis pfefferi* | (320 - 29th ; 650 - 99th) | 38 |
| | | | | | | | | *Pseudosimochromis curvifrons* | (275 - 28th ; 600 - 99th) | 28 |
| | *A. calliptera* (Masoko) | | | | | | | Benthic | (3800 - 21st; 5700 - 98th) | 70 |
| | | | | | | | | Littoral | (3800 - 21st; 5700 - 98th) | 69 |
| Killifish | Killifish | *Nothobranchius furzeri* | GCA_001465895.2 | | 15816766 | 1200 | 2.6E-09 | Chefu | (275 - 39th ; 620 - 99th) | 42 |
| | | | | | | | | Limpopo | | 41 |
| Clownfish | Clownfish | *Amphiprion clarkii* | GCA_027123335.1 | | 27144647 | 846 | 4.00E-08 | Papua New Guinea | (800 - 10th; 1600 -99th) | 48 |
| | | | | | | | | Indonesia | | 37 |
| Sticklebacks | Sticklebacks | *Gasterosteus aculeatus* | GCA_016920845.1 | | 13025426 | 472 | 5.05E-09 | Finger Lake | (4000 - 8000) | 35 |
| | | | | | | | | Long Lake | | 48 |
| | | | | | | | | Spirit Lake | | 44 |
| | | | | | | | | Watson Lake | | 47 |

**Table S2:** This table gives supplementary information to Figure 3C. The estimates, p-value, and their levels of significance, with '.'<0.1; '*' <0.05; '**' <0.005; and '***' <0.0005, are reported for each species/population.

| **Species** | **Estimate** | **p-value** | **signif. level** |
|---|---|---|---|
| *J. pfefferi* | 1.22E-14 | 0.9 | |
| *P. curvifrons* | -1.53E-13 | 0.08 | . |
| *P. nyererei* | -8.05E-14 | 0.036 | * |
| *A. nubila* | -9.29E-15 | 0.4848 | |
| *A. calliptera* – benthic | -9.22E-14 | 0.02169 | * |
| *A. calliptera* – littoral | -3.74E-14 | 0.08 | . |
| *N. furzeri* – Chefu | -6.41E-15 | 0.33 | |
| *N. furzeri* – Limpopo | -2.14E-14 | 0.04 | * |
| *A. clarkii* – Papua New Guinea | -6.89E-13 | 0.003 | ** |
| A. clarkii – Indonesia | -1.61E-12 | 0.0006 | *** |
| *G. aculeatus* – Finger Lake | -2.78E-14 | 0.03 | * |
| *G. aculeatus* – Long Lake | -1.32E-13 | 0.003 | ** |
| *G. aculeatus* – Watson Lake | -1.22E-13 | 0.002 | ** |
| *G. aculeatus* – Spirit Lake | -1.19E-13 | 0.0003731 | *** |

**Table S3:** Population recombination divergence index values. For each comparison, we reported the $d_{me}$, $d_{ms}$, PRDI, and the associated 95% confidence interval calculated across the whole genome.

| Group | Comparison | | Dme | Dms | PRDI | **95% CI** |
|---|---|---|---|---|---|---|
| | Population 1 | Population 2 | | | | |
| Victoria cichlids | *Pundamilia nyererei* | *Astatotilapia nubila* | 0.0407 | 0.011 | 0.0297 | [0.017;0.050] |
| Tanganyika cichlids | *Jabarichromis pfefferi* | *Pseudosimochromis curvifrons* | 0.37 | 0.0722 | 0.297 | [0.255;0.340] |
| *A. calliptera* (Masoko) | Benthic | Littoral | 0.148 | 0.046 | 0.102 | [0.078;0.118] |
| Killifish | Limpopo | Chefu | 0.134 | 0.0334 | 0.101 | [0.076;0.127] |
| Clownfish | Papua New Guinea | Indonesia | 0.167 | 0.0596 | 0.107 | [0.094;0.124] |
| Sticklebacks | Finger lake (Benthic) | Long Lake (Limnetic) | 0.168 | 0.0736 | 0.0941 | [0.072;0.125] |
| | Watson Lake (Benthic) | Spirit Lake (Limnetic) | 0.0866 | 0.0281 | 0.0584 | [0.049;0.080] |

**Table S4: PRDI per chromosome.** PRDI values and 95% confidence intervals for each chromosome and each comparison. Chromosome names are those of the corresponding assembly. PRDI values significantly different than zero are in bold.

| Cichlids | | | | | | | *G. aculeatus* | | | | | *N. furzeri* | | | *A. clarkii* | | |
|---|---|---|---|---|---|---|---|---|---|---|---|---|---|---|---|---|---|
| | *A. calliptera* | | Vic. Cichlids | | Tan. Cichlids | | | Finger - Long | | Spirit - Watson | | | | | | | |
| **Chr. Name** | **PRDI** | **CI 95 %** | **PRDI** | **CI 95 %** | **PRDI** | **CI 95 %** | **Chr. Name** | **PRDI** | **CI 95 %** | **PRDI** | **CI 95 %** | **Chr. Name** | **PRDI** | **CI 95 %** | **Chr. Name** | **PRDI** | **CI 95 %** |
| LS420019.2 | **0.212** | [0.131;0.293] | 0.029 | [-0.050;0.095] | **0.199** | [0.012;0.376] | NC_053212.1 | 0.060 | [-0.001;0.105] | **0.050** | [0.027;0.083] | NC_029649.1 | **0.103** | [0.035;0.147] | CM049567.1 | **0.140** | [0.069;0.177] |
| LS420020.2 | 0.035 | [-0.028;0.132] | 0.034 | [-0.035;0.068] | **0.206** | [0.052;0.335] | NC_053213.1 | 0.025 | [-0.074;0.176] | **0.034** | [0.011;0.104] | NC_029650.1 | 0.062 | [-0.037;0.126] | CM049568.1 | **0.172** | [0.087;0.241] |
| LS420021.2 | 0.066 | [-0.009;0.144] | 0.046 | [-0.010;0.089] | **0.364** | [0.250;0.432] | NC_053214.1 | 0.088 | [-0.014;0.297] | **0.044** | [0.001;0.093] | NC_029651.1 | 0.043 | [-0.062;0.130] | CM049569.1 | **0.068** | [0.014;0.212] |
| LS420022.2 | -0.006 | [-0.101;0.101] | -0.021 | [-0.110;0.046] | **0.377** | [0.156;1.030] | NC_053215.1 | 0.087 | [-0.009;0.188] | **0.049** | [0.016;0.144] | NC_029652.1 | **0.221** | [0.111;0.307] | CM049570.1 | **0.123** | [0.082;0.229] |
| LS420023.2 | **0.121** | [0.042;0.181] | 0.014 | [-0.049;0.066] | **0.425** | [0.160;0.651] | NC_053216.1 | 0.062 | [-0.005;0.086] | 0.033 | [-0.033;0.082] | NC_029653.1 | 0.008 | [-0.081;0.109] | CM049571.1 | **0.029** | [0.002;0.082] |
| LS420024.2 | **0.103** | [0.034;0.143] | 0.052 | [-0.020;0.099] | **0.288** | [0.032;0.406] | NC_053217.1 | **0.122** | [0.083;0.214] | **0.042** | [0.028;0.103] | NC_029654.1 | 0.121 | [-0.024;0.146] | CM049572.1 | **0.095** | [0.046;0.128] |
| LS420025.2 | **0.145** | [0.020;0.226] | 0.050 | [-0.019;0.139] | **0.254** | [0.173;0.314] | NC_053218.1 | **0.044** | [0.024;0.208] | **0.120** | [0.069;0.215] | NC_029655.1 | 0.064 | [-0.030;0.179] | CM049573.1 | **0.166** | [0.091;0.283] |
| LS420026.2 | 0.049 | [-0.081;0.238] | 0.037 | [-0.012;0.106] | **0.142** | [0.063;0.258] | NC_053219.1 | **0.085** | [0.052;0.168] | **0.061** | [0.008;0.080] | NC_029656.1 | 0.004 | [-0.098;0.133] | CM049574.1 | **0.086** | [0.000;0.144] |
| LS420027.2 | **0.112** | [0.036;0.154] | **0.065** | [0.011;0.138] | **0.232** | [0.127;0.273] | NC_053220.1 | 0.136 | [-0.010;0.161] | **0.050** | [0.009;0.102] | NC_029657.1 | 0.114 | [-0.003;0.153] | CM049575.1 | **0.096** | [0.020;0.177] |
| LS420028.2 | **0.092** | [0.008;0.119] | -0.026 | [-0.084;0.064] | **0.373** | [0.235;0.494] | NC_053221.1 | **0.073** | [0.014;0.180] | **0.063** | [0.042;0.102] | NC_029658.1 | **0.093** | [0.037;0.178] | CM049576.1 | **0.098** | [0.035;0.156] |
| LS420029.2 | **0.137** | [0.012;0.310] | 0.032 | [-0.058;0.096] | **0.351** | [0.142;0.422] | NC_053222.1 | 0.059 | [0.037;0.205] | **0.083** | [0.053;0.123] | NC_029659.1 | 0.014 | [-0.033;0.150] | CM049577.1 | **0.083** | [0.040;0.097] |
| LS420030.2 | **0.101** | [0.009;0.187] | **0.092** | [0.035;0.133] | **0.233** | [0.137;0.292] | NC_053223.1 | **0.103** | [0.031;0.275] | **0.065** | [0.018;0.099] | NC_029660.1 | 0.057 | [-0.005;0.131] | CM049578.1 | **0.078** | [0.022;0.167] |
| LS420031.2 | 0.092 | [-0.049;0.158] | 0.053 | [-0.013;0.133] | **0.184** | [0.073;0.393] | NC_053224.1 | 0.081 | [-0.092;0.151] | **0.062** | [0.023;0.079] | NC_029661.1 | 0.100 | [-0.099;0.185] | CM049579.1 | **0.086** | [0.043;0.133] |
| LS420032.2 | 0.172 | [-0.005;0.217] | -0.042 | [-0.116;0.035] | **0.257** | [0.182;0.400] | NC_053225.1 | **0.140** | [0.030;0.223] | **0.085** | [0.031;0.097] | NC_029662.1 | 0.103 | [-0.044;0.140] | CM049580.1 | **0.155** | [0.062;0.229] |
| LS420033.2 | 0.021 | [-0.142;0.106] | 0.067 | [-0.057;0.171] | **0.194** | [0.094;0.323] | NC_053226.1 | 0.030 | [-0.048;0.101] | **0.047** | [0.029;0.089] | NC_029663.1 | -0.036 | [-0.115;0.198] | CM049581.1 | **0.047** | [0.005;0.104] |
| LS420034.2 | 0.075 | [-0.074;0.217] | 0.085 | [-0.014;0.099] | **0.194** | [0.107;0.287] | NC_053227.1 | **0.100** | [0.076;0.207] | **0.043** | [0.030;0.153] | NC_029664.1 | 0.099 | [-0.079;0.148] | CM049582.1 | **0.106** | [0.027;0.179] |
| LS420035.2 | **0.096** | [0.020;0.142] | 0.065 | [-0.040;0.088] | **0.201** | [0.047;0.240] | NC_053228.1 | 0.059 | [-0.009;0.091] | 0.072 | [-0.023;0.114] | NC_029665.1 | 0.094 | [-0.056;0.128] | CM049583.1 | **0.080** | [0.002;0.195] |
| LS420036.2 | 0.100 | [-0.012;0.207] | 0.098 | [-0.042;0.139] | **0.172** | [0.060;0.390] | NC_053229.1 | -0.054 | [-0.127;0.030] | **0.038** | [0.024;0.049] | NC_029666.1 | 0.052 | [-0.054;0.168] | CM049584.1 | **0.188** | [0.095;0.218] |
| LS420037.2 | -0.002 | [-0.132;0.156] | 0.042 | [-0.007;0.194] | **0.415** | [0.236;0.719] | NC_053231.1 | **0.134** | [0.076;0.295] | **0.077** | [0.036;0.112] | NC_029667.1 | **0.129** | [0.024;0.231] | CM049585.1 | **0.069** | [0.036;0.195] |
| LS420038.2 | 0.072 | [-0.041;0.204] | 0.034 | [-0.107;0.084] | **0.459** | [0.208;0.924] | NC_053232.1 | **0.130** | [0.032;0.151] | **0.081** | [0.015;0.146] | | | | CM049586.1 | **0.101** | [0.066;0.141] |
| LS420039.2 | **0.122** | [0.015;0.287] | 0.090 | [-0.215;0.155] | **0.460** | [0.136;0.688] | | | | | | | | | CM049587.1 | **0.119** | [0.043;0.135] |
| LS420040.2 | **0.194** | [0.063;0.225] | 0.038 | [-0.012;0.100] | **0.271** | [0.048;0.436] | | | | | | | | | CM049588.1 | **0.059** | [0.014;0.185] |
| | | | | | | | | | | | | | | | CM049589.1 | **0.121** | [0.060;0.163] |
| | | | | | | | | | | | | | | | CM049590.1 | **0.196** | [0.109;0.479] |
